# Prediction of phase separation propensities of disordered proteins from sequence

**DOI:** 10.1101/2024.06.03.597109

**Authors:** Sören von Bülow, Giulio Tesei, Kresten Lindorff-Larsen

## Abstract

Phase separation is thought to be one possible mechanism governing the selective cellular enrichment of biomolecular constituents for processes such as transcriptional activation, mRNA regulation, and immune signaling. Phase separation is mediated by multivalent interactions of biological macromolecules including intrinsically disordered proteins and regions (IDRs). Despite considerable advances in experiments, theory and simulations, the prediction of the thermodynamics of IDR phase behaviour remains challenging. We combined coarse-grained molecular dynamics simulations and active learning to develop a fast and accurate machine learning model to predict the free energy and saturation concentration for phase separation directly from sequence. We validate the model using both experimental and computational data. We apply our model to all 27,663 IDRs of chain length up to 800 residues in the human proteome and find that 1,420 of these (5%) are predicted to undergo homotypic phase separation with transfer free energies *<* −2*k*_B_*T*. We use our model to understand the relationship between single-chain compaction and phase separation, and find that changes from charge-to hydrophobicity-mediated interactions can break the symmetry between intra-and inter-molecular interactions. We also analyse the structural preferences at condensate interfaces and find substantial heterogeneity that is determined by the same sequence properties as phase separation. Our work refines the established rules governing the relationships between sequence features and phase separation propensities, and our prediction models will be useful for interpreting and designing cellular experiments on the role of phase separation, and for the design of IDRs with specific phase separation propensities.

## Introduction

Biomolecular condensates are large, dynamic assemblies of macromolecules in the cell. In contrast to membrane-bound organelles, biomolecular condensates are not enclosed by a lipid bilayer and their composition is predominantly governed by the differences in inter-molecular interactions between macromolecules inside and outside the condensate, and with the solvent.^1–3^ Various types of these condensates have been identified in the cell, including the nucleolus, Cajal bodies, and promyelocytic leukaemia (PML) bodies in the nucleus; and P bodies, stress granules, and Balbiani bodies in the cytosol.^1,3,4^ Physiological functions of biomolecular condensates include buffering of local and cellular concentrations, response to stimuli and stress, transcriptional regulation, or gating through the nuclear pore complex, among others.^2,5^

The biophysical origins of condensate formation in the cell are an active area of research. Phase separation (PS) coupled to percolation, involving the reversible de-mixing of solutes into biomolecule-dense and dilute phases, is thought to be one of the mechanisms underlying condensate formation. ^6^ Multivalent interactions, often involving intrinsically disordered regions (IDRs) of proteins, contribute to driving PS.^7,8^ Polymer theory has proven to be a powerful foundation to interpret *in vitro* experiments on PS, and has been particularly useful for understanding the phase behaviour of a single species of IDRs. ^5,7,9,10^ In practice, PS of IDRs depends on the IDR sequence and external conditions such as temperature and the type and concentration of ions in solution.^1,11^

Experiments, theory and simulations have been used together to shed light on the rules governing PS *in vitro* and *in vivo*. The sticker-and-spacer model has proven successful in rationalizing and predicting sequence-dependent PS.^7,12^ When this framework is applied to IDRs, amino acid residues are categorized into stickers, which contribute the major driving force for PS through for example hydrophobic, *π*-*π*, and electrostatic interactions; and spacers, which intersperse the stickers and contribute weaker interactions. The patterning of sticker residues along the linear sequence determines the condensate-spanning network of sticker-sticker interactions and, thereby, the extent of de-mixing. On the other hand, spacers influence the solubility of the macromolecules and modulate PS propensities and, thereby, the extent of de-mixing.^6^

A number of previous studies have helped uncover how sequence properties of IDRs affect PS. Through the design of constructs of different repeat sequences, Quiroz & Chilkoti tuned the upper and lower critical solution temperatures (UCST and LCST, respectively) of synthetic IDRs and proposed a set of sequence rules governing PS, including molecular weight, zwitterionic character, aromaticity, and arginine content.^13^ A number of subsequent studies have further highlighted the important roles of aromatic residues as stickers^9,12^ and ranked them in the order Phe<Tyr<Trp based on their relative strength in driving PS.^9,12,14–16^ Studies on different IDRs have also identified Arg as a sticker, primarily thought to be due to its interactions with aromatic residues, although Arg–Arg interactions may also play a role,^17^ while Lys has been characterized as a spacer.^12,14,18^ Moreover, PS is also affected by substitutions between spacer residues, such as Gly-to-Ser, Gly-to-Ala, Ser-to-Thr, and Asn-to-Gln. ^18,19^ The effects are governed by changes in the solvation volume of the IDR and are sensitive to sequence context and solution conditions. ^18,19^ Both aromatic and charge patterning have been shown to measurably influence PS of IDRs;^9,20–22^ thus even at fixed amino acid composition, the sequence patterning may affect PS substantially as shown for example by shuffled variants of the low complexity domain (LCD) of hnRNPA1^22^, the LAF-1 RGG domain^14^ and NICD.^21^

Coarse-grained simulations of physics-based models of IDRs with residue-level resolution have been instrumental in elucidating the sequence dependence of PS. ^9,15,23–29^ In many of these models, short-ranged interactions are described using a modified Lennard-Jones (LJ) potential, where the stickiness of each residue is captured by amino acid-specific parameters. Additionally, salt-screened electrostatic interactions between charged residues are described using the Debye-Hückel potential.^23,24^ Investigations of PS using these models commonly employ direct coexistence simulations, wherein a single condensate is formed in an elongated simulation box, making it pseudo-infinite along the shorter box sides and thereby reducing finite-size effects.^23^ Dignon et al. combined single-chain and direct coexistence simulations of the LCD of FUS, hnRNPA2, LAF1, TDP-43, and their variants to investigate the relationship between IDR single-chain expansion and multi-chain PS. ^10^ The authors found a strong correlation between UCST and the Θ temperature at which the isolated IDR has ideal-chain compaction, i.e., a Flory scaling exponent *ν* of 0.5. Further, applying analytical theory to sequences composed of equal numbers of Lys and Glu, Lin & Chan showed that UCST vs. radius of gyration (*R*_g_) follows a power law whereas UCST depends linearly on sequence parameters quantifying charge patterning,^30^ i.e., sequence charge decoration (SCD)^31^ and *κ*.^32^ This coupling between single-chain compaction and propensity to undergo PS has been exploited to develop transferable residue-level models through data-driven approaches in which models are trained on data from experiments probing single-chain conformational properties.^9,15,24–27,29,33^ We have developed one such model, CALVADOS, by deriving the stickiness parameters from experimental small angle X-ray scattering and paramagnetic relaxation enhancement NMR data.^15,27^ These physics-based models accurately estimate the propensity of IDRs of diverse sequences to undergo PS^18,24,27,28^ and capture the decoupling between single-chain compaction and PS propensities for sequences with large absolute values of the net charge per residue (NCPR).^18,27^

Polymer theory and coarse-grained simulations of IDRs have also highlighted a strong correlation between the second virial coefficient and PS propensity, ^10,34^ which led to the development of an analytical model for predicting PS of IDR mixtures based on two-body IDR–IDR interactions.^35^ The relationship between chain compaction, virial coefficients and PS have also led to approaches to use single-chain simulations to predict phase diagrams for IDRs.^36,37^ Strategies to quantify interactions in phase-separating IDR mixtures have also been developed.^7,38^ Recent work identified connections between the second virial coefficient, mobility, and PS propensity of IDRs.^39^ The authors used an active learning protocol to learn and characterize the trade-off between PS propensity and protein mobility in condensates from coarse-grained simulations,^39^ and to define molecular features to generate solutions of multiple components in distinct phases of different composition.^40^

Sequence-based predictors of PS behaviour *in vitro* and *in vivo* have been developed, either employing heuristic rules or based on supervised machine-learning (ML) approaches. The ML predictors are often trained on experimental data (e.g., *in vivo* PS databases) to learn sequence rules governing *in vitro* PS propensities or *in vivo* localization to biomolecular condensates. These predictors generally aim to classify IDRs into two groups—phase separating and not phase separating—and estimate the probability to undergo PS of a given IDR without quantifying transfer free energy or saturation concentration. For example, DeePhase is trained using sequence feature embeddings to distinguish PS-prone IDR sequences from structured proteins and non-PS-prone IDRs.^41^ FuzDrop predicts the droplet-promoting propensity of proteins based on the entropy differences in the bound and unbound state.^42^ The catGRANULE algorithm predicts a granule-localization propensity from sequence using features including RNA binding and structural disorder propensities. ^43^ PScore predicts PS propensity from *π*-*π* interaction frequencies alone.^44^ PSAP and ParSe (v2) are classifiers trained on curated *in vitro* and *in vivo* PS databases to predict if a protein undergoes PS based on sequence features.^45,46^ FINCHES uses parameters from coarse-grained force fields and a mean-field approach to estimate homo- and heterotypic interactions including semiquantitative estimates of phase diagrams.^47^ Simulations have also been used to derive rules enabling predictions of variations in PS of specific families of IDRs. ^9,48^

Despite the many advances highlighted above, it is still challenging to accurately predict the concentrations of the dense and, importantly, the dilute phase, even for *in vitro* systems of a single species of IDR in solution. In turn, predicting the free energy of transfer from dilute into the dense phase is likewise challenging, in particular due to the sensitivity of the dilute phase (saturation) concentration to sequence changes.^18,19^

Here, we exploit the accuracy of coarse-grained simulations to estimate the PS propensity of IDRs^15,28,33,38,49^ to develop a machine learning model that efficiently predicts phase behaviour of single-component protein solutions from sequence across a broad region of sequence space. As the reference physics-based model we use CALVADOS, which recently enabled the characterization of structural ensembles of all IDRs in the human proteome, i.e., the human IDRome.^50^ While simulations of single chains are extremely fast, simulating a system of ≈ 100 chains using CALVADOS requires on the order of several days on a modern GPU. Therefore, a simulation screen of PS propensities for the whole human IDRome would be computationally extremely expensive.

To overcome this limitation, we here develop and employ an active learning protocol to select ≈400 sequences for direct-coexistence simulations with CALVADOS. We then use the results to train a neural network regression model that accurately predicts saturation concentrations and transfer free energies of single-component protein solutions *in vitro* directly from sequence. Through extensive validation against both simulation and experimental data, we show that our machine learning model has an accuracy on par with CALVADOS simulations at a fraction of the computational cost, and use the results to shed light on the interplay between sequence features that determine homotypic PS. Finally, we exploit the wealth of simulation data to study structural properties of the condensates and their interfaces.

## Results and Discussion

### An active learning protocol to predict transfer free energies from sequence

We have previously shown that CALVADOS simulations give rise to dilute phase concentrations that are in good agreement with experiments for a range of proteins and variants.^27,27^ We therefore aimed to develop a PS predictor based on results from phase coexistence simulations using CALVADOS 2^15,27^ (Fig. 1A). We used an active learning protocol to generate a diverse set of training data covering a large feature space in the human IDRome, so as to allow the model to correlate a broad range of sequence features to PS propensities (Fig. 1A). For the purpose of the active learning protocol, we initially built a support vector regression (SVR) model to predict the propensity of IDR systems to undergo homotypic PS, expressed as transfer free energies Δ*G*,

**Figure 1.**
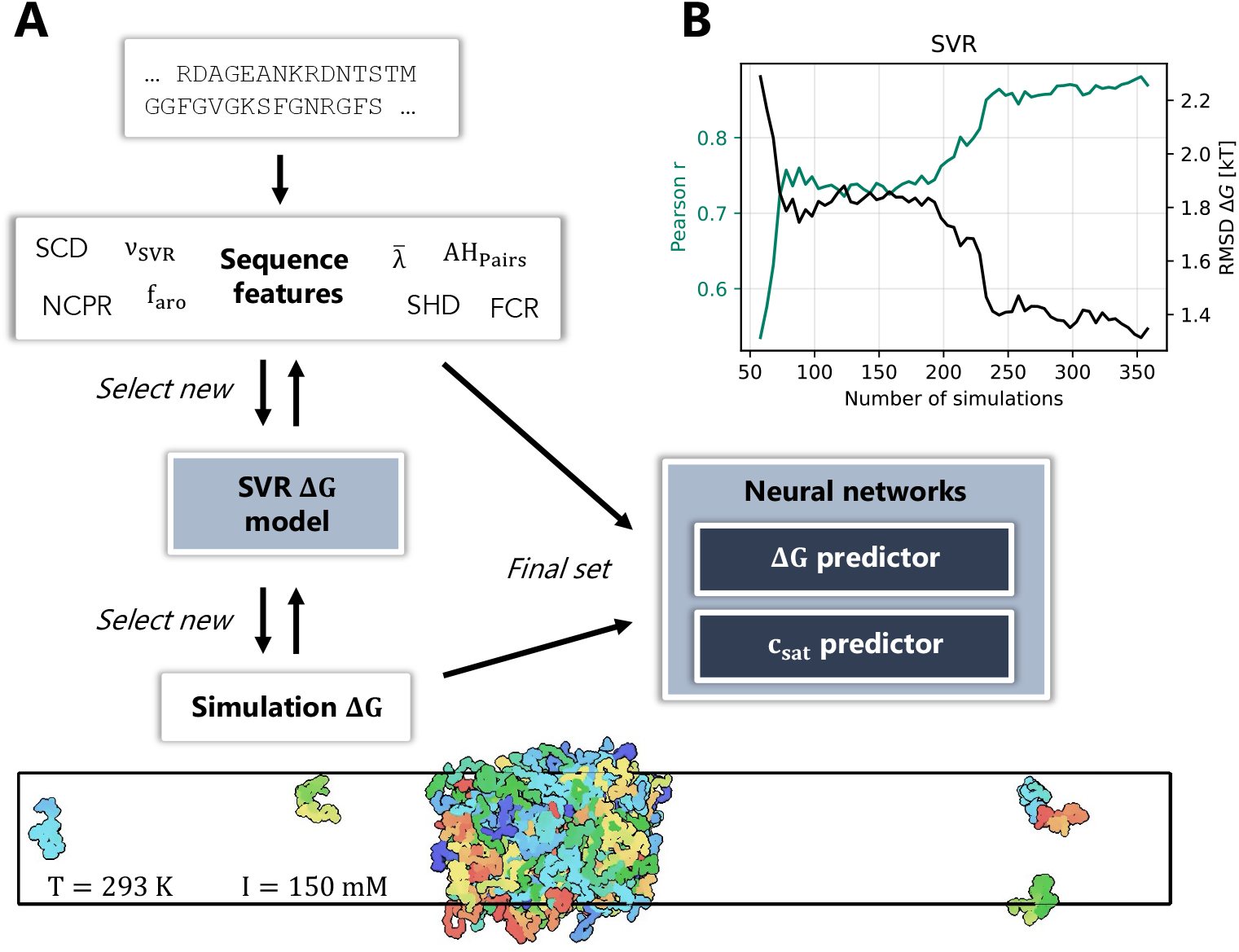
An active learning framework for predicting phase separation. (A) Active-learning protocol to train a PS predictor from simulation data. SVR: Support vector regression. The iterative sampling and training was driven by SVR models; once sampling had been completed we trained dense neural networks to predict the Δ*G* values and the saturation concentrations. (B) Convergence of the active learning protocol for an IDRome_90_ validation set (n=27).

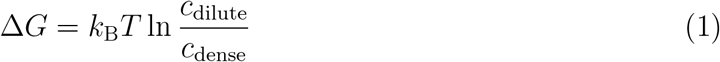

from sequence features (see Methods). We fixed simulation conditions to *T* = 293 K and ionic strength of *I* = 150 mM to be compatible with many *in vitro* experiments. The dynamic range of the simulations is roughly Δ*G* = −10 *k*_B_*T* to Δ*G* ≈ 0 *k*_B_*T* ; sequences that give rise to Δ*G <* −10 *k*_B_*T* have so few proteins in the dilute phase during the simulation time that they cannot be distinguished. Similarly, sequences that are not predicted to undergo spontaneous PS (Δ*G >* 0 *k*_B_*T*) will be assigned to Δ*G* = 0 *k*_B_*T* because we cannot detect any stable condensate (see Methods).

At each learning iteration, we re-trained the SVR model on the current set of Δ*G* values collected from the coexistence simulation results to predict Δ*G* from sequence input features. The input features encode the physics of the CALVADOS 2 force field (Table S1. The model selected new sequences to simulate out of a pool of 90% of the human IDRome (IDRome_90_). The remaining 10% (IDRome_10_) were held out as a validation set that we only examined after having finalized model development and training. Briefly described, our active learning protocol selected new sequences for simulation based on three conditions: (1) Large range of Δ*G* values (roughly Δ*G* ≈ −10 *k*_B_*T* to Δ*G* ≈ 0 *k*_B_*T*), (2) highest inter-model uncertainty in cross-validation, so as to select new sequences that the model was unsure about, and (3) large coverage of input sequence feature space.

We monitored the convergence of the active learning protocol by calculating Pearson’s correlation coefficients (*r*) and root-mean-squared deviations (RMSD) between the SVR predictions and simulations of Δ*G* via cross-correlation (80% training, 20% test) as a function of the number of simulation sequence data points used for training (Fig. S1A). The values of RMSD and *r* reached a plateau beyond ≈ 250 simulations (with a total of 362 simulations), and scatter plots of predicted vs. simulated Δ*G* show that the model can distinguish different PS propensities (Fig. S1B). We therefore tested the convergence of the model for a set of 27 independent sequences from IDRome_90_ (Fig. 1B). We observed a strong improvement of the prediction accuracy up to ≈ 250 included simulated sequences, with only small improvements beyond. We therefore concluded that the training has converged.

### Dense neural network improves prediction accuracy and is transferable

Having established convergence of the SVR prediction model, we pooled all simulation data from training and convergence test within the IDRome_90_ set, resulting in 362 + 27 = 389 sequences. We used these data to train two slightly different dense neural networks (NN): The first model predicts the transfer free energy Δ*G* (Eq. 1), whereas the second model predicts the natural logarithm of the saturation mass concentration of the IDR, i.e. of the dilute phase concentration in the coexistence simulations. We optimized the network architecture via a grid search in parameter space (Fig. S2). Different architectures with two hidden layers gave very similar prediction performance (as measured by RMSD). We selected the model with hyperparameters *α* = 5 and 2 × 10 hidden layers for its combination of high performance and speed.

The resulting Δ*G* and ln *c*_sat_ models showed excellent prediction accuracy, as measured by cross-validation (Fig 2A,D). To test if the NN models can be generalized to previously unseen data (i.e. data outside the sequence pool that could be selected during active learning of the SVR model), we predicted Δ*G* values for 26 held-out IDRome_10_ sequences. We find that the models predict Δ*G* and ln *c*_sat_ for these independent sequences as accurately as for the IDRome_90_ sequences, thus concluding that the models predict Δ*G* and ln *c*_sat_ with *r >* 0.9 and RMSD< 1 (Fig. 2B,E).

**Figure 2.**
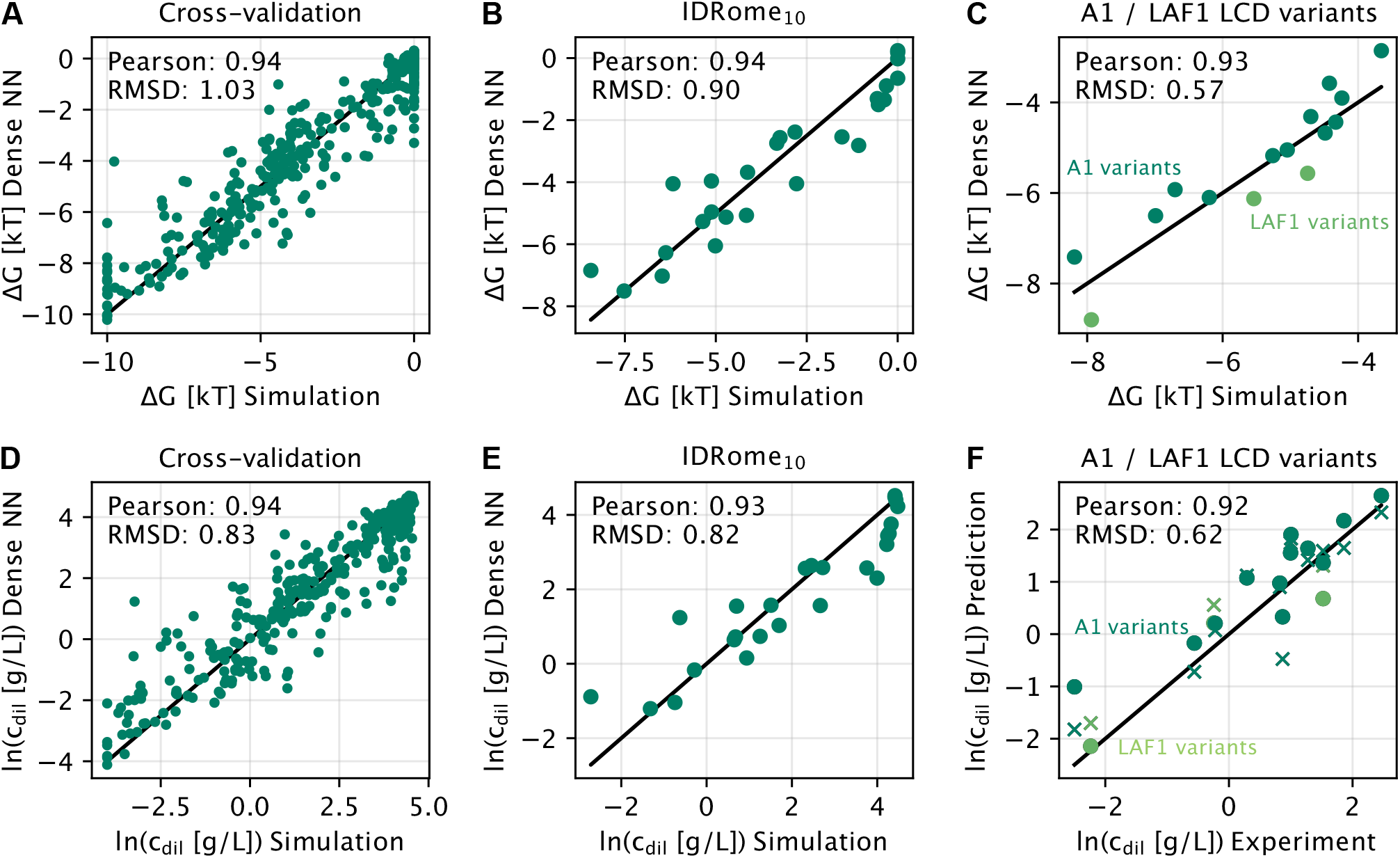
Accurate machine learning models enable quantitative predictions of phase separation. Results from (A, D) IDRome_90_ cross-validation, (B, E) IDRome_10_ validation, and (C, F) further simulation/experimental validation for NN predictors of Δ*G* (A-C) and the natural logarithm of the saturation concentration (D–F). Circles in (F) represent predictions by the NN and crosses represent CALVADOS 2 simulation results.

### Benchmarking the prediction model with experimental data

The good prediction of CALVADOS 2 Δ*G* values by the NN model is encouraging, as the CALVADOS 2 model in turn has been fine-tuned to match experimental saturation concentrations.^15^ We therefore aimed to directly compare the NN predictions with experimental data. We collected simulation and experimental PS data of the LCD of hnRNPA1 and LAF1, as well as variants thereof, from the original CALVADOS 2 parameterization work, none of which were used during training of the NN models.^15^ Remarkably, sequence variant effects for both the simulation Δ*G* values (Fig. 2C) and the experimental saturation concentrations (Fig. 2F) were predicted very accurately by the NN model, with RMSD= 0.62 and Pearson’s *r* = 0.92, on par with the simulation results (RMSD= 0.60, *r* = 0.91).

### PS predictions are interpretable with sequence features

We used the NN model to predict Δ*G* for all sequences in the human IDRome, again noting the dynamic range of our simulations and analyses corresponds to −10 ≲ Δ*G/k*_B_*T* ≲ 0. The distribution of Δ*G* is strongly skewed towards sequences with weak or non-PS values (Fig. S3). Only 571 (2%), 892 (3%), or 1,420 (5%) out of the 27,663 sequences in the IDRome (≤ 800 residues) are predicted to undergo PS when using PS thresholds of Δ*G <* −4 *k*_B_*T*, -3 *k*_B_*T*, or -2 *k*_B_*T*, respectively. Therefore, only a small fraction of IDRs in the IDRome are predicted to undergo PS without partners at the given conditions (*T* = 293 K, *I* = 150 mM).

We correlated the predicted Δ*G* values with each of the individual sequence features that we use as input to the NN model (Fig. 3). As expected, the mean sequence hydrophobicity, hydrophobic patterning, charge patterning, and predicted single-chain scaling exponent all correlate positively with increased predicted PS propensity (low Δ*G*). Lower absolute NCPR likewise correlates with lower Δ*G*. Thus, the NN learned overall effects of physical properties that have previously been shown to affect PS and which are captured in the CALVADOS model. The high standard deviations across individual bins indicate that none of the individual features we analysed can quantitatively predict the PS propensities. In addition to highlighting the complex interplay between features, we also note that some of the features have been derived to capture properties of the sequence at fixed composition and length, and were therefore not designed to be used alone across the diverse set of IDRome sequences.^51^

**Figure 3.**
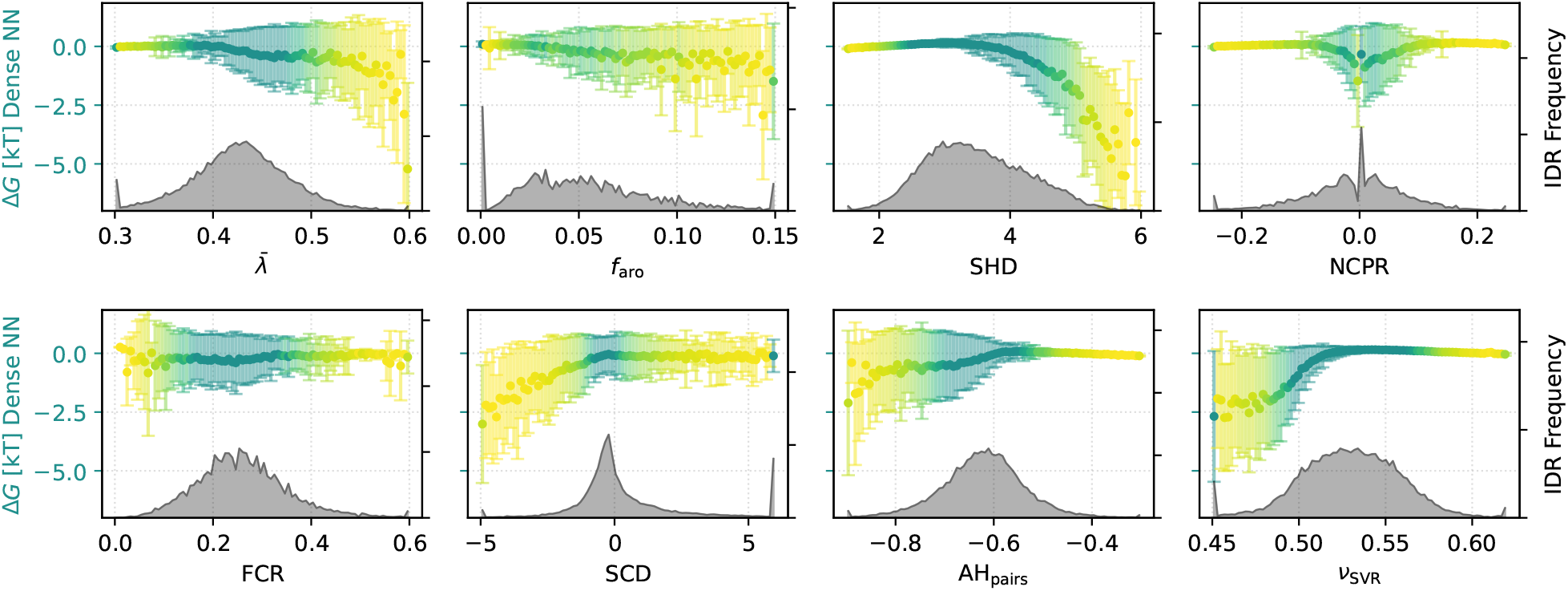
Correlation of IDRome Δ*G* with input sequence features. Error bars indicate the standard deviation per bin. Grey lower histograms indicate the number of sequences per bin. Colours indicate number of sequences per bin corresponding to the histograms below, with darker colour indicating more proteins.

We also investigated the dependence of Δ*G* on two combined features, e.g., 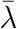 and all other features, or SCD and all other features (Figs. S4 and S5). The corresponding 2D histograms show which combinations of features allow a clear distinction between low and high Δ*G* values. The combinations 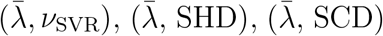 show clear Δ*G* separation potential, as do the combinations (SCD, FCR), (SCD, SHD), and (SCD, *ν*_SVR_). Like our previously described model for single chain compaction,^50^ our model therefore likely uses several related features to disentangle effects of sequence composition, patterning and length.

We also trained Δ*G* NN models on a reduced set of input features, using only one feature or combinations of two features as input (Fig. S6). Combinations of descriptors of sequences hydrophobicity and single-chain scaling expansion (which itself uses several features as input) performed best. All models using one or two input features were much less predictive compared to the full model with all features, necessitating the full model for quantitative predictions of Δ*G*.

We investigated, how confidently the model predicts Δ*G* for different regions in feature space. Using the data from 389 simulation results in the IDRome_90_ set, we trained a model to predict the unsigned prediction error of Δ*G* (Fig. S7) based on sequence features and predicted Δ*G* values. The prediction error model underestimated the compounded simulation and prediction error (RMSD = 0.7 *k*_B_*T* vs. 1.0 *k*_B_*T*) and is only weakly correlated with the true absolute difference of Δ*G* and predicted Δ*G* (*r* = 0.54; Fig. S7A). The error for the Δ*G* model and the predicted error for the Δ*G* model only depend weakly on the simulated Δ*G* (Fig. S7B,C). In light of these results, we instead report the RMSD of the IDRome_10_ validation set (RMSD(Δ*G*)=0.90 *k*_B_*T* and RMSD(ln *c*_sat_)=0.82) as global estimates of the prediction errors.

### Correlation between single-chain features and PS propensity

Previously, the relationship between sequence, single chain features and PS propensities have been studied. In particular, it has been shown that measures of single-chain compaction such as the Flory scaling exponent, *ν*, are correlated with the PS propensity for related variants of given sequences. ^9,10,30^ We leveraged our fast model to screen thousands of sequence variants in order to learn which features might affect *ν* and Δ*G* differently. To this aim, we performed Monte Carlo (MC) sampling in sequence space to explore how our Δ*G* model reacts to sequence perturbations, starting from a range of weakly to intermediately PS-prone sequences (−4 < Δ*G/k*_B_*T <* −1).

We first determined the effect of free sequence exploration on Δ*G* via swap moves, i.e., reshuffling the residues of a given IDR composition (Fig. S8). We observed clear positive correlations between changes in *ν*_SVR_ and Δ*G* as well as SCD and Δ*G*, in agreement with earlier findings.^10,30^ In contrast, we do not see a strong effect of hydrophobic patchiness (SHD) for a given composition.

We then asked, which changes in sequence features might possibly break the correlation between single-chain scaling exponent *ν*_SVR_ and PS propensity, i.e., which changes in the sequence reduce or increase Δ*G* while maintaining fixed single-chain expansion *ν*. We therefore performed a MC walk in sequence space towards low predicted Δ*G*.

We first restricted the MC algorithm to only swap moves while restraining *ν*_SVR_ close to their original values. Given these restraints, Δ*G* values could barely move away from their starting values (Fig. S9). The patchiness of charges and hydrophobic residues increased with PS propensity and *ν*_SVR_ until *ν*_SVR_ reached the pre-set restraint tolerance, beyond which the MC algorithm was stuck, with overall absolute changes in Δ*G <* 0.4 *k*_B_*T*.

We modified the algorithm in a second step, now allowing substitutions to any of the 19 other residue types (i.e. changing sequence composition) to assess, how *ν*_SVR_ and PS propensity are globally decoupled. We fixed NCPR alongside *ν*_SVR_ in this step, as we expected the effect of net charge to dominate more subtle effects.^11,18,21,27^ During the MC walks towards low Δ*G* with swap and substitution moves, sequences increased in hydrophobicity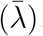, whereas the fraction of charged residues decreased to maintain the same single-chain compaction (*ν*) (Fig. 4). Thus we find that, for fixed single-chain compaction, hydrophobic sequences tend to phase separate more strongly than sequences whose compaction is driven by charge interactions.

**Figure 4.**
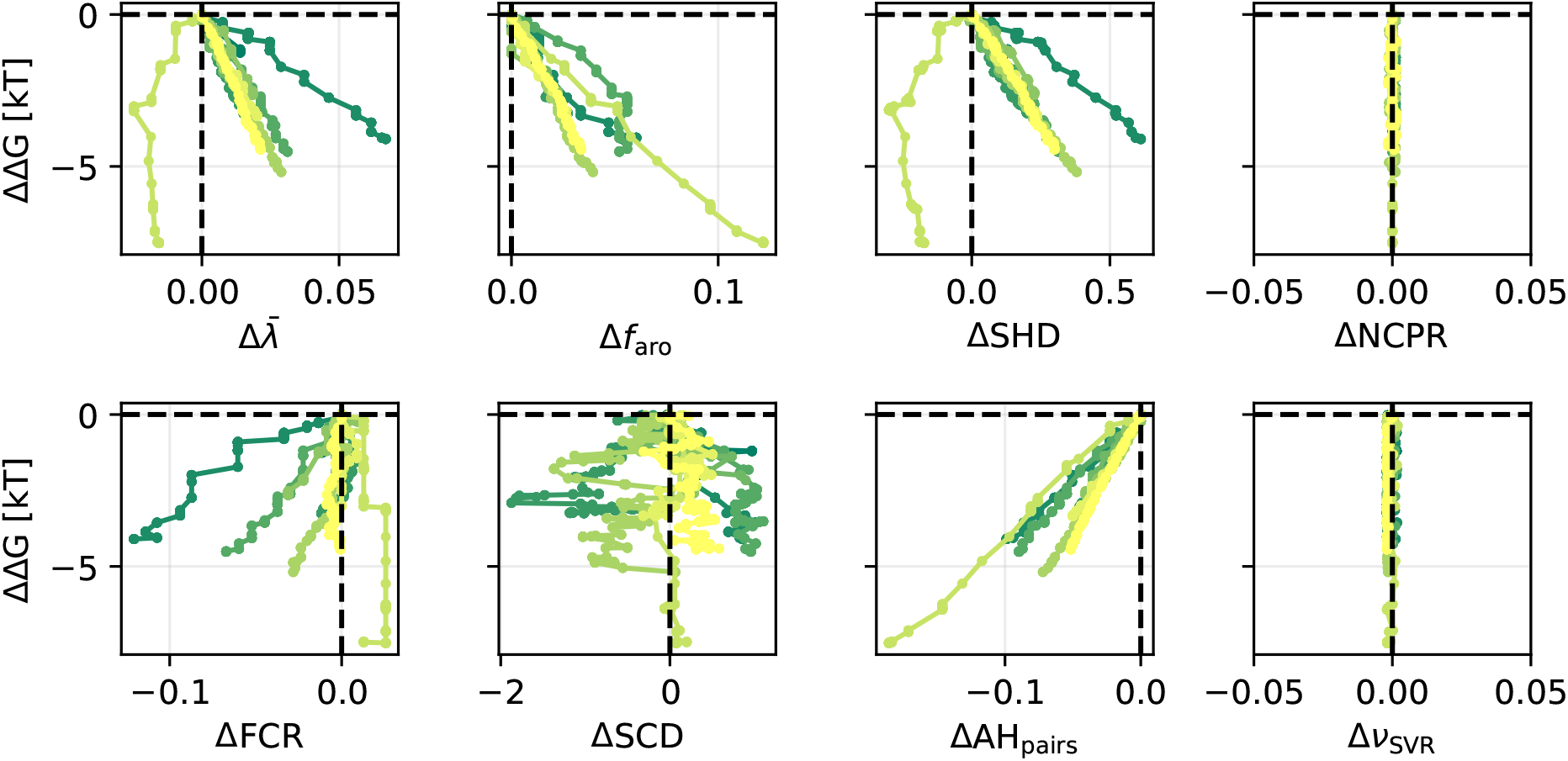
Effect of Monte-Carlo sequence optimization towards strong PS (target Δ*G* = −10 *k*_B_*T*) starting from random sequences in window −4 < Δ*G/k*_B_*T <* −1 using swap moves and single substitutions while restraining *ν*_SVR_ and NCPR. Different colours correspond to independent runs of the algorithm.

The key findings from our sequence exploration runs are: (1) For a given sequence composition, changes in SCD and *ν*_SVR_ are strongly correlated with changes in PS propensities. (2) For a given composition, Δ*G* and *ν*_SVR_ are so tightly coupled, that we could not substantially move one without the other. (3) Globally, hydrophobic sequences with low charge content (and low patterning) as well as less hydrophobic sequences with higher charge content can have the same *ν*_SVR_ but substantially different PS propensities, with the former showing stronger PS.

### Variations in structural properties at the condensate interface

In line with expectations for homopolymers, we and others have previously found that IDRs are more expanded in homotypic condensates than in dilute solution of a poor solvent (water).^18,27,36^ To examine these effects more broadly, we calculated *ν* in the dilute and dense phases of 110 of the 389 training data sequences that we simulated during the active learning protocol and which had −10 < Δ*G/k*_B_*T <* −4. While the chain compaction in the dilute phase varies substantially across sequences, in agreement with the compaction estimated from single-chain simulations, the IDRs all have *ν* ≈ 0.5 (Fig. S10) in the dense phase, in line with the condensates acting as a Θ solvent for the IDRs.

The substantial variation and differences in structural properties in dilute and dense phases suggest that there might also be variation in structural properties at the condensate interfaces. Farag et al.^33^ used lattice simulations to examine the structural preferences, chain expansion and orientation of the LCD of hnRNPA1 in the dense phase, condensate interface, and dilute phase, and found both increased chain expansion and a propensity to take on an orientation perpendicular to the interface for chains located at the droplet interface. In other studies—using different simulation frameworks, analysis methods and IDR sequences—chains at the interface have been found to be more compact than in the dense phase.^52–54^

We used our large-scale direct coexistence simulations of substantially different IDR sequences to quantify structural preferences at the interface and compare them to those in the dilute and dense phase. We calculated bin-weighted^33^ profiles of the radius of gyration (*R*_g_) along the direction normal to the condensate interface (that is along the *z* axis) from the 110 direct coexistence simulations (Fig. 5A and additional examples in Fig. S11). To compare the compaction across sequences, we normalized the *R*_g_(*z*) values by the average value in the dense phase. In line with the calculations of *ν* (Fig. S10), we find that the expansion at the interface is generally between that in the dilute and dense phase (Fig. 5B). In line with previous findings,^33^ we find, however, substantial complexity in the structural properties along the interface for many sequences (see Fig. S11 for examples), and for some sequences we find that parts of the interface have a bin-weighted *R*_g_ greater than the dense phase (Fig. 5B).

**Figure 5.**
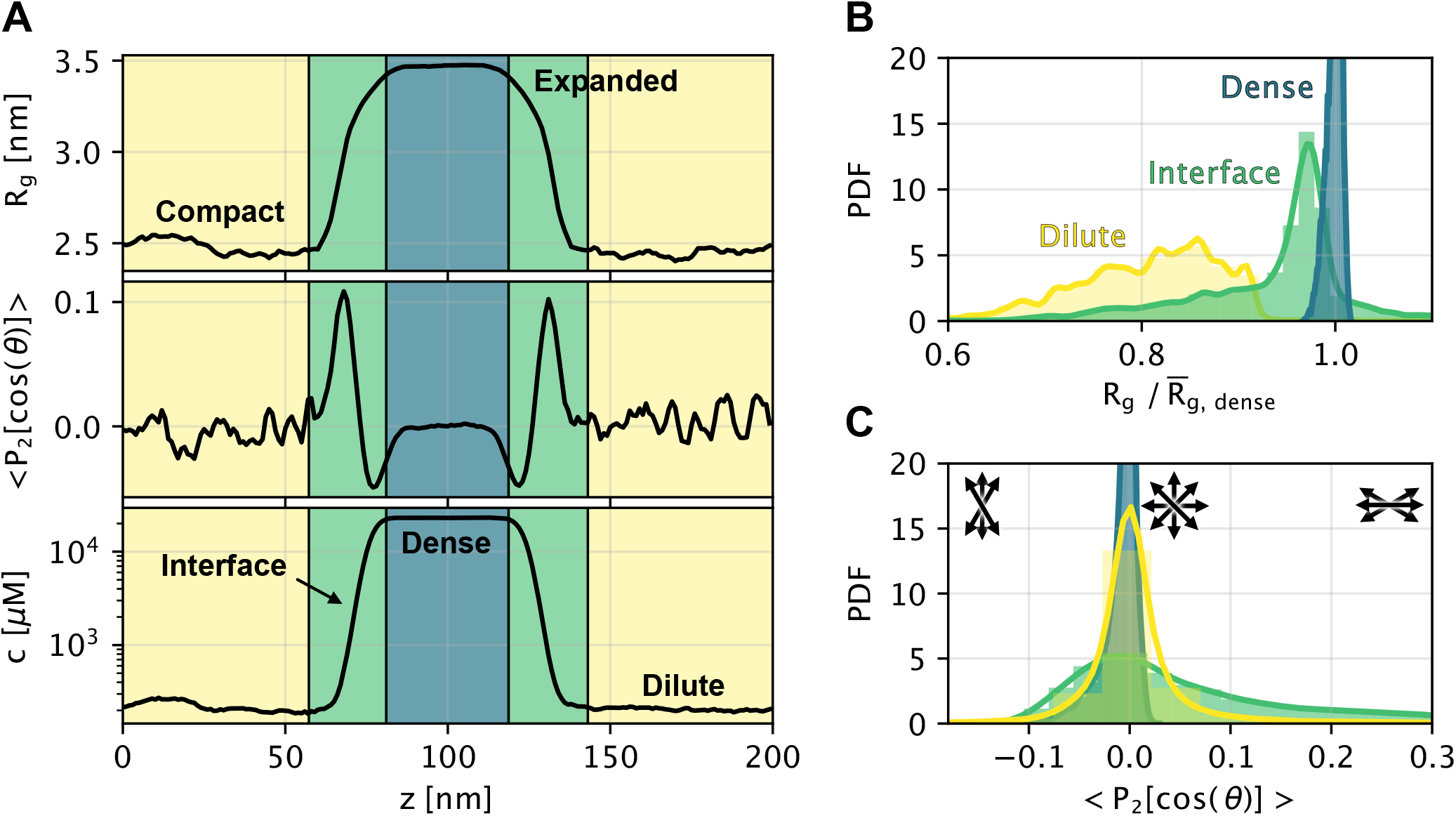
Structural properties in the dilute phase, interface region and dense phase. (A) Example of profiles of *R*_g_(*z*), orientation (*S*_*z*_(*z*) = ⟨ *P*_2_[cos(*θ*)] ⟩), and protein concentration (*c*(*z*)) for bins along the long box edge *z*. Coloured shading indicates the dilute phase (yellow), interface (green), and dense phase (blue) regions. (B) Histograms of 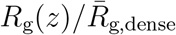 distributions from the bin values for all proteins separately for bins in the dense phase, interface, and dilute phase. All bins contribute equally to these distributions, regardless of chain or monomer concentration per bin (Methods). (C) Histograms of *S*_*z*_ distributions from pooled bin values for all proteins, as for (B). Black arrows illustrate preferential orientations, whereby *S*_*z*_ = 0 corresponds to an isotropic (random) orientation, *S*_*z*_ *>* 0 indicates preferential orientation along *z* (normal to the interface), and *S*_*z*_ < 0 indicates preferential orientation orthogonal to *z* (along the interface). We assign small deviations from *S*_*z*_ = 0 in the dilute phase to be statistical noise from the low amount protein in the dilute phase of the most strongly phase separating proteins.

Inspired by previous analyses of IDR orientation,^33,52,55^ we calculated a chain order parameter, *S*_*z*_, to quantify the extent to which chains are aligned along the *z*-axis. *S*_*z*_ = 1 corresponds to full alignment along the *z*-axis (normal to the interface), an isotropic distribution of orientations gives *S*_*z*_ = 0, and *S*_*z*_ = −1*/*2 indicates alignment orthogonal to *z*, i.e. parallel to the condensate surface. We calculated *S*_*z*_ for each chain and time step and averaged *S*_*z*_ values for each bin along the *z*-axis to obtain an orientation profile along *z*. As expected, we find close-to-random orientations in both the dilute and dense phases (Figs. 5A, 5B and S11). In contrast, we find much greater variation in the behaviour at and near the interfaces, with many sequences showing both positive and negative peaks of *S*_*z*_ in the interface regions (Figs. 5B and S11). In many cases we find *S*_*z*_ < 0 closest to the dense phase and *S*_*z*_ *>* 0 further out in the interface region. In line with findings for the hnRNPA1 LCD,^33^ we find that the IDRs in the interface region have a preference to be oriented perpendicularly to the interface.

Having found considerable variation in the level of compaction and orientational preferences in the interface region, we asked whether these differences were correlated with sequence and structural features of the IDRs. We find a strong correlation between 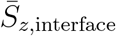 and 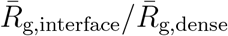 so that those sequences that are most expanded at the interface are also those that have the strongest preference to be oriented perpendicularly to the interface (Fig. S12). We also find that these values are both correlated with Δ*G*, so that the sequences with the strongest driving force for PS also show largest values of 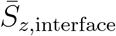 and 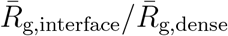(Fig. S12). Since Δ*G* is correlated with several sequence features (Figs. 3, S4 and S5), these features also correlate with the structural preferences in the interface region.

### Limitations

CALVADOS was trained to reproduce biophysical measurements of single-chain conformational properties, and has been shown to reproduce both single-chain and PS properties it was not trained on. We therefore rationalized that we could build an accurate prediction method for PS by targeting CALVADOS simulations. Nevertheless, these *in silico* predictions of homotypic PS may not capture all relevant properties of the densely crowded, heterogeneous environment in the cell.^56–59^ For example, while CALVADOS has been shown to capture effects of varying the ionic strength on PS,^15^ it will not capture specific effects due to ion-specific asymmetrical partitioning in condensates.^60^ Similarly, sequences that do not undergo homotypic PS (for example highly charged sequences) may undergo PS with oppositely charged molecules in the cell. Likewise, other discrepancies between the *in vitro* and *in vivo* conditions will limit the model. While we have validated our prediction methods for natural sequences from the human IDRome, it is possible that they will be less accurate for non-natural sequences. We note, however, that sequence design based on CALVADOS has shown transferability outside the realm of natural sequences.^22^

Furthermore, our predictors inherit the strengths and limitations of the CALVADOS 2 model. In particular, the Δ*G* and *c*_sat_ estimations from CALVADOS 2 direct coexistence simulations have an absolute relative error, ⟨|*c*_sat, sim_ −*c*_sat, exp_|*/c*_sat, exp_⟩, of 90%,^61^ corresponding to a RMSD of ln(*c*_sat_[g/L]) of 0.73. We deliberately trained our model to reproduce PS at a fixed set of temperature and ionic strength. Even though it could be retrained at different conditions, the CALVADOS model does not fully capture variation of PS with temperature, as only the electrostatic term of the force field is temperature dependent via Eq. 5, whereas the effect of temperature on residue stickiness is not captured in the model. Furthermore, the description of electrostatic interactions based on a Debye-Hückel screening term with fixed cutoff of 4 nm is limited both for very high and low ionic strengths as well as ion-type and pH-specific effects.^11,60,62^

## Conclusion

We have developed machine learning models to quantitatively predict homotypic PS of IDRs at physiologically relevant conditions. We devised and implemented an active learning approach to select the most relevant simulation data to train a model that estimates PS globally across diverse sequences. While previous models have been developed to classify sequences into those that PS and those that do not, we are not aware of other models to predict the saturation concentration and transfer free energies for a wide set of disordered proteins sequences.

Since PS may be a generic property of a wide range of proteins^63,64^ and cellular protein concentrations can vary substantially, we envisage that the quantitative aspect our model will be particularly important; because many proteins may undergo PS at some concentration it is not always clear which conditions a binary PS prediction method refers to. Our results are thus complementary to exciting new work by Ginell et al. ^47^, published as a preprint alongside this manuscript. Leveraging the pairwise interaction parameters of CALVADOS 2^15,27^ and a modified form of Mpipi^28,65^ in a mean-field approach, the authors developed a model to rapidly compute interaction maps and semi-quantitative phase diagrams between any pair of disordered proteins, validating their method with a range of biologically interesting systems.

Condensate interfaces have unique chemical properties and are thought to play potential roles in both function and pathology.^7^ We have analysed structural features of IDRs in the dilute and dense phases, as well as the important and unique interface region, and correlated these with the sequences of the IDRs. We find substantial variation in the conformational properties at interfaces that can be explained by the same features that drive formation of condensates. We also find substantial fine structure and heterogeneity at the interfaces, and future work is aimed towards understanding the molecular origins of these effects.

We envisage that our prediction methods may become valuable tools for experimentalists and theoreticians to obtain rapid and accurate estimates of *in vitro* PS propensities of IDRs before performing costly experiments or simulations, and to design and interpret experimental and computational studies. Our machine learning models may also be used to explore more widely the relationship between sequence and PS properties and to link biological properties, disease and PS. The code for our model is freely available, and we also provide easy access via an online implementation as a Google Colab notebook. Finally, by providing access to a unique and large set of direct-coexistence simulations for a wide range of sequences, we enable detailed analysis and insights into the relationship between sequence and PS properties including analyses of the structure and thermodynamics of PS.

## Methods

### CALVADOS 2 force field

We performed molecular dynamics simulations using the coarse-grained CALVADOS 2 model.^15^ As with similar HPS models, ^23,66^ each protein residue is represented by one bead with size *σ* and interaction strength *λ*.

The full model is a linear combination of contributions to the potential energy,

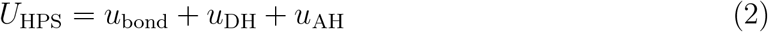

with *u*_bond_ the bonded potential, *u*_DH_ a Debye-Hückel electrostatic potential, and *u*_AH_ is the Ashbaugh-Hatch modification of a Lennard-Jones potential.^66^

Beads of neighbouring residues in the sequence are connected by bonds described by a harmonic potential,

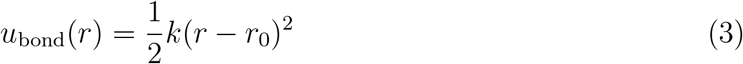

using *k* = 8033 kJmol^*−*1^nm^*−*2^ as force constant and *r*_0_ = 0.38 nm as equilibrium distance.

A Debye-Hückel potential describes the solvent-screened electrostatic interactions,

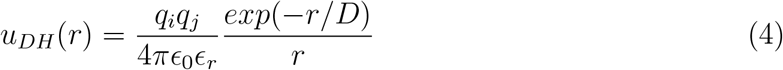

with *q*_*i*_ the charge of bead *i, ϵ*_0_ the vacuum permittivity, 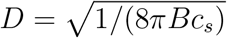 the Debye length of an electrolyte solution of ionic strength *c*_*s*_, and *B*(*ϵ*_*r*_) the Bjerrum length of temperature-dependent dielectric constant *ϵ*_*r*_,^67^

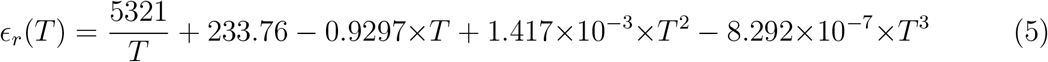

Electrostatic interactions were truncated and shifted at the cutoff distance *r*_*c*_ = 4 nm.

Nonelectrostatic nonbonded interactions were represented by a truncated and shifted Ashbaugh-Hatch (AH) potential^66^. It is a scaled Lennard-Jones (LJ) potential of the following functional form,

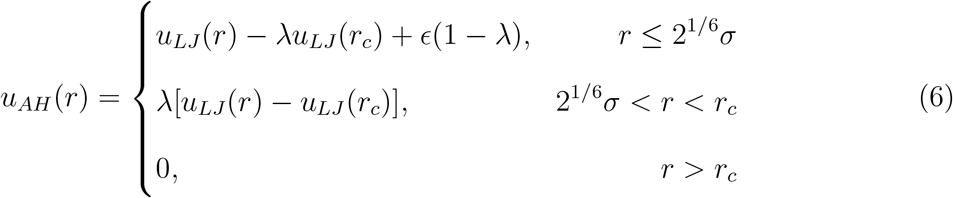

with *σ* = (*σ*_*i*_ + *σ*_*j*_)*/*2, *λ* = (*λ*_*i*_ + *λ*_*j*_)*/*2 for residues *i* and *j*, and the LJ potential

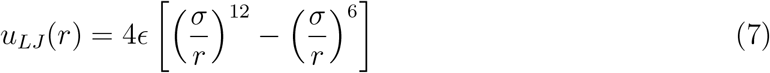

where *ϵ* = 0.8368 kJ mol^*−*1^ and *r*_*c*_ = 2 nm.

### Molecular dynamics simulations and estimation of PS propensities

We used the openMM v8.0 simulation package^68^ to perform molecular dynamics simulations. Proteins were inserted into an elongated simulation box with dimensions 25 nm × 25 nm × 300 nm for sequences with more than 350 residues and 20 nm × 20 nm × 200 nm otherwise. Initial configurations were fully elongated proteins (along the z direction) packed in parallel in the box centre in the *z* direction.

We performed simulations in the NVT ensemble with a Langevin integrator (*γ* = 0.01 ps^*−*1^) with timestep of 0.01 ps. Protein configurations were saved at either 1 ns or 10 ns intervals. The first 600 ns of each simulation were discarded to account for equilibration. ^15^ All simulations were run at temperature *T* = 293 K and ionic strength *I* = 150 mM.

Time-averaged concentrations in the dense and dilute phases (particle density maps) were calculated with custom scripts. ^69^ The scripts center the slab in the *z* direction based on a heuristic estimate of the centre of density. This analysis assumes that there is at most one condensed phase in the simulation. Visual inspection of all density time series revealed that 12 simulations from the IDRome_90_ training set and 2 simulations from the IDRome_10_ validation set showed the presence of two or more condensed phases; these likely represent simulations that did not converge to a single stable phase during the pre-defined simulation time. Since the analysis framework would erroneously interpret a smaller condensate as belonging to the dilute phase, and thus overestimate *c*_sat_, these were removed from model training and validation. We note that the two simulations that were removed from the IDRome_10_ set were identified before assessing model accuracy. We show the time series for the 14 simulations in Fig. S13.

In order to compare chain expansion from slab and single chain simulations, we performed single chain simulations for a subset (*n* = 110) of sequences simulated in the course of the active training protocol that show PS with −10 < Δ*G/k*_B_*T <* −4. We performed each single chain simulation in a simulation box of 25 nm × 25 nm × 25 nm at the same ensemble and conditions as the direct coexistence simulations. Simulations were carried out for 200 ns simulation time. The first 20 ns of each simulation run were discarded as equilibration.

The boundaries between dense and dilute phase were determined by fitting a hyperbolic tangent to the concentration profile, as described previously:^15^

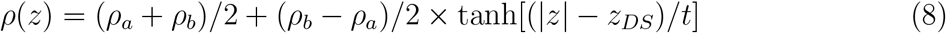

with *ρ*_*a*_ and *ρ*_*b*_ the densities of the dense and dilute phases, respectively.

The dense and dilute phases are estimated to be in regions |*z*| *< z*_*DS*_ − *β*_dense_*t* and |*z*| *> z*_*DS*_ + *β*_dilute_*t*, with *β*_dense_ = 1.5 and *β*_dilute_ = 2.5 (for sequences A8K8P3 740 1157, O94906 1 81, Q96SB4 1 59, Q8N9I0 83 138, Q4V348 1 281, O15504 1 116, Q86W67 1 206, Q9BWV2 1 254) or *β*_dilute_ = 5 otherwise. Here, *z*_*DS*_ and *t* are the position of the dividing surface and thickness of the interface, respectively. We defined the interface as the zone between the dense and dilute phase, i.e., the region *z*_*DS*_ − *β*_dense_*t <* |*z*| *< z*_*DS*_ + *β*_dilute_*t*.

### IDR sequence selection by active learning

We selected IDR sequences for phase coexistence simulations in a multi-step process that we devised to maximize model performance at minimal computational cost. Before initiating the model we selected 10% of the IDRome (IDRome_10_) to be used for final assessment of the model and did not analyse these sequences until the final analysis.^70^ The remaining 90% of the IDRome are denoted as IDRome_90_.

We first collected initial seed simulations performed at the same temperature and ionic strength in previous work. The seed consisted of 38 YTH domain protein IDRs^69,71^ and 28 additional simulations from unpublished projects.

We then devised an active learning protocol to explore new IDR sequences for simulation. During each step in the active learning procedure, we trained a new support-vector regression (SVR) model with parameters *C* = 10 and *ϵ* = 0.01 to predict transfer free energies Δ*G* (Eq. 1 to partition into the dense phase. We used the sklearn python package^72^ for all ML models in this work. Out of a pool of sequence features, the algorithm selects the combination of three features that gives the best prediction (measured by the Pearson correlation coefficient, *r*)The pool of features consisted of *N*, 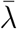, *f*_aro_, SHD, NCPR, FCR, SCD, *κ, R*_g_, *ν*, and *M*_w_. The features are defined in Table S1. In these analyses we used *ν* obtained from analyses of single chain simulations.^50^ We determined the prediction accuracy as the average Pearson correlation coefficient, *r*, on the validation set from 50 cross-validations for each feature combination, each with 80% and 20% of simulations randomly chosen as training and validation set, respectively. The set of 50 models with average highest-performing input feature combination was then used to predict Δ*G* for all sequences in the IDRome_90_ set, resulting in 50 predicted Δ*G* per IDRome_90_ sequence. Under the assumption that a large cross-model uncertainty indicates lack of accuracy for specific types of sequences in the IDRome_90_,^70^ we restricted the pool of new sequences to simulate to the top 100 sequences with highest Δ*G* variance. Out of these 100 sequences, we picked 5–10 sequences maximizing the distance in feature space, as calculated by the Mahalanobis distance (*d*_M_). We first selected the highest variance sequence for simulations; then we selected a second sequence (out of the 100 sequences) with the highest *d*_M_ to the first sequence, then a third sequence by maximizing the sum of *d*_M_ to the first two sequences etc., resulting in 5–10 new sequences to simulate based on available computational resources at each iteration. Based on this protocol, we iteratively selected and simulated a total of 137 sequences.

Following this first phase of sequence exploration, we modified the active learning algorithm to focus the learning on a more uniform range of predicted Δ*G* values. We therefore added another criterion to the procedure in the above described protocol: In the modified protocol, we selected the top 5–10 sequences with highest cross-model variances (top 50%) and *d*_M_ separately for bins of Δ*G* (in units of *k*_B_*T*): [−∞, −6], [−6, −5], [−5, −4], [−4, −3], [−3, −2], [−2, −1], and [−1, 1]. In this way, we selected sequences with different values of predicted Δ*G* for further simulations. We iteratively selected and simulated 179 additional sequences based on this modified protocol.

Once the model appeared to have converged, we selected additional sequences for a final convergence test from within the IDRome_90_ set, drawing 4 new sequences randomly from each predicted Δ*G* bin, with same brackets as above.

### Dense neural models to predict transfer free energies and saturation concentrations from the final set of simulations

We built and trained two small dense neural networks (NN) to predict Δ*G* and ln *c*_sat_ from sequence features. These models were trained on the final set of 389 phase coexistence simulations gathered from the three-step procedure described above.

We chose the input features listed in Table S1 except *N*, *M*_*w*_, *κ*, as those showed limited prediction accuracy alone or using pairs of features (Fig. S6). We also removed *R*_*g*_ to restrict the input to features that can rapidly be generated from sequence without requiring simulation work. In the SVR models described above we used values for the Flory scaling exponent (*ν*) based on single chain simulations;^50^ for the NN we instead used an accurate sequence-based SVR model *ν*_SVR_.^50^ The prediction of *ν*_SVR_ in turn uses SCD, SHD, *κ*, FCR, and 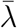 as input features.^50^ The final input features for the NN were thus 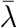, *f*_aro_, SHD, NCPR, FCR, SCD, AH_pairs_, and *ν*_SVR_. We note that several of these features were designed to be used individually for fixed sequence composition and length, and that combining them as input to the NN helps overcome this limitation. The AH_pairs_ is a new feature that we designed for this work to quantify the interaction between chains. For each residue pair in the protein, AH_pairs_ calculates a score based on the *u*_AH_ term for hydrophobic interactions in Eq. 2 scaled by the interaction volume (Table S1).

We performed a hyperparameter grid optimization for *α* and architecture of hidden layers, converging on a final set of parameters, *α* = 5 and two hidden layers of 10 nodes each (Fig. S2). As for the SVR model, the accuracy of the model was determined by 50 cross-validations (80% training, 20% validation), using Pearson’s *r* and RMSD as metrics.

We selected 26 sequences from the IDRome_10_ for final assessment of model accuracy. As for the IDRome_90_ convergence test above, the sequences were selected randomly from bins of predicted Δ*G* values, now using the NN predictor instead of the SVR predictor to sort sequences into Δ*G* bins. We used the same Δ*G* bin definitions as above.

### Monte-Carlo simulations in sequence space

We performed Monte-Carlo (MC) sampling in sequence space to explore how sequence variations by swaps or substitutions relate to changes in PS propensities. The sequence length *N* was fixed to the initial sequence length.

At each iteration, the algorithm chose randomly between swap moves or substitutions with equal probability (unless only swap moves were allowed). If swap moves were chosen by the algorithm, the residue types of 10 pairs of positions in the sequence were swapped (attempted swaps of identical residue types or positions led to repeated tries). If substitution moves were chosen by the algorithm, 10 random residues along the sequence were substituted with any of the other 19 residue types with equal probability.

The algorithm computed the features and predicted Δ*G* of the resulting trial sequences. The set of 10 moves (swap or substitutions) were collectively accepted or rejected by the algorithm. To be accepted, the features needed to be within tolerance of the constraints, where applicable (*ν* tolerance: 0.001, NCPR tolerance: 0.002). In addition, the predicted Δ*G* of the trial sequence needed to satisfy a Metropolis criterion,

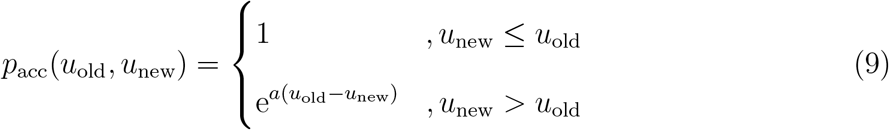

with *a* = 100 and *u* = *k*(*x* − *x*_*t*_)^2^, where *x* and *x*_*t*_ are current and target value, respectively, and *k* = 0.3.

### Analysis of structural properties in condensates

We calculated the Flory scaling exponent *ν* separately for the dense phase, interface, and dilute phase. We root-mean-square (RMS) averaged all intra-protein residue distances 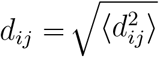 (for pairs of residues *i, j* separated with sequence distance |*j* − *i*|) from proteins with centre-of-mass in the designated region (e.g. dilute phase). *ν* was then obtained from a fit of *d*_*ij*_ = *R*_0_|*j* − *i*|^*ν*^ to the data, with *R*_0_ as flexible fit parameter and |*j* − *i*| *>* 5.

In order to compute binned profiles of *R*_*g*_ vs. *z*-position, we computed 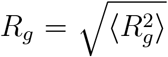 for all chains and trajectory frames. We constructed histograms of *R*_*g*_(*z*) by distributing the calculated chain *R*_*g*_ values to the *z*-positions of the residue beads of the protein, following the method in Farag et al. ^33^. For each bin, we then calculated an RMS-averaged *R*_*g*_.

We calculated an order parameter (*S*_*z*_) to quantify the extent to which chains are aligned along the *z*-axis:

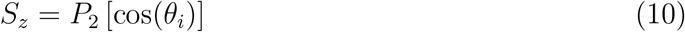

Here, 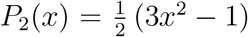 is the second Legendre polynomial, and *θ*_*i*_ the angle between the smallest principal axis of the chain *i* (corresponding to longest chain elongation) and the *z*-axis ([0,0,1]) of the simulation box. As for *R*_*g*_ above, we calculated *S*_*z*_ for every protein chain at each time frame, and performed bin-wise averaging along *z* using the *z*-positions for each amino acid residue (bead) in the protein, resulting in a single mean ⟨*S*_*z*_⟩ value.

In this way, the *z*-dependent profiles represent the average *R*_*g*_ and *S*_*z*_ of all frames and chains while accounting for the inhomogeneous distribution of protein bead positions for each IDR.^33^

## Data and code availability

Data and code used for this work is available via https://github.com/KULL-Centre/_2024_buelow_PSpred. An web implementation of the neural network models can also be run using https://colab.research.google.com/github/KULL-Centre/_2024_buelow_PSpred/lob/main/PSLab.ipynb. Our simulation data is available via https://sid.erda.dk/sharelink/hlZfnFz4AM.

## Acknowledgements

We thank Alex Holehouse, Tanja Mittag, and Thea Klarsø Schulze for helpful discussions and suggestions. We also thank Alex Holehouse for his willingness to delay preprinting of their manuscript such that both preprints could reference one another. S.v.B acknowledges support by the European Molecular Biology Organisation through Postdoctoral Fellowship grant ALTF 810-2022. K.L.-L. acknowledges support by the Novo Nordisk Founda-tion via the PRISM (Protein Interactions and Stability in Medicine and Genomics) centre (NNF18OC0033950). We acknowledge access to computational resources from the Biocomputing Core Facility at the Department of Biology, University of Copenhagen, from the Resource for Biomolecular Simulations (ROBUST; supported by the Novo Nordisk Foundation; NNF18OC0032608), and the Danish National Supercomputer for Life Sciences (Computerome).

## Supporting Information

**Table S1:** Sequence features used in this work.

|  |  |
| --- | --- |
| $N$ | Sequence length. |
| $\bar{\lambda}$ | Mean sequence hydrophobicity $\bar{\lambda} = \frac{1}{N} \sum_{i=1}^N \lambda_i$ for sequence of length $N$. We use the $\lambda$ values from the CALVADOS 2 model.<sup>15</sup> |
| $f_{\text{aro}}$ | Fraction of aromatic residues, $f_{\text{aro}} = \frac{1}{N} \sum_{i=1}^N a_i$ with $a_i = 1$ for residues Phe, Tyr, Trp, and $a_i = 0$ otherwise. |
| SHD | Sequence hydropathy decoration<sup>73</sup> using $\lambda$ values from CALVADOS 2. |
| NCPR | Net charge per residue, $\text{NCPR} = \frac{1}{N} \sum_{i=1}^N q_i$ with $q_i$ the charge per residue. The N-terminus and C-terminus are positively and negatively charged, respectively. |
| FCR | Fraction of charged residues $\text{FCR} = \frac{1}{N} \sum_{i=1}^N Q_i$. $Q_i = 1$ for nonzero charges, $Q_i = 0$ otherwise. |
| SCD | Sequence charge decoration.<sup>31</sup> |
| $\text{AH}_{\text{pairs}}$ | $\text{AH}_{\text{pairs}} = \frac{1}{N(N+1)/2} \sum_{i=1}^N \sum_{j=1}^N \int_{2^{1/6}\sigma}^{r_c} 4\pi r^2 u_{\text{AH}}^{i,j}$. Sum of scaled attractive part of integrated pair potential for all pairs of sequence residues. $u_{\text{AH}}$ corresponds to the Ashbaugh-Hatch potential Eq. 6. |
| $\nu_{\text{SVR}}$ | SVR model for Flory scaling exponent.<sup>50</sup> |
| $\kappa$ | Charge patterning parameter.<sup>32</sup> |
| $R_g$ | Radius of gyration in nm. |
| $M_w$ | Molecular weight in Dalton. |

**Figure S1:**
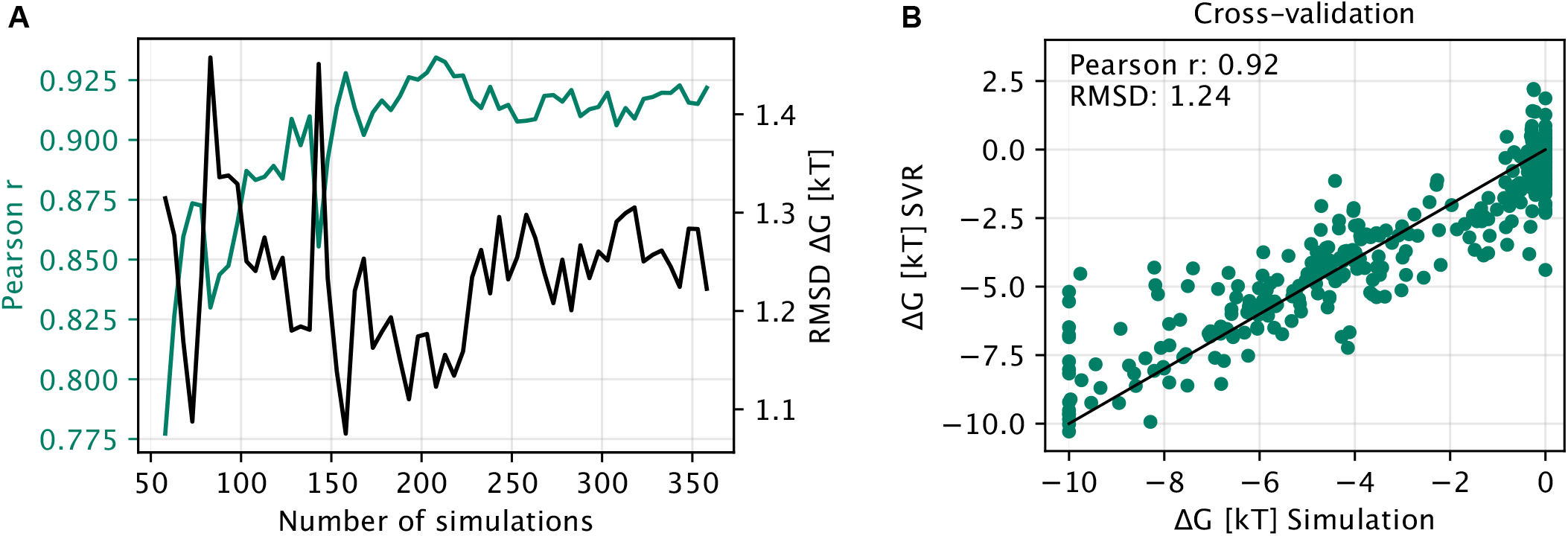
(A) SVR model cross-validation Pearson *r* and RMSD of prediction of Δ*G* for increasing numbers of simulated sequences. Scatter plot of simulated vs. SVR predicted Δ*G* values (n=362).

**Figure S2:**
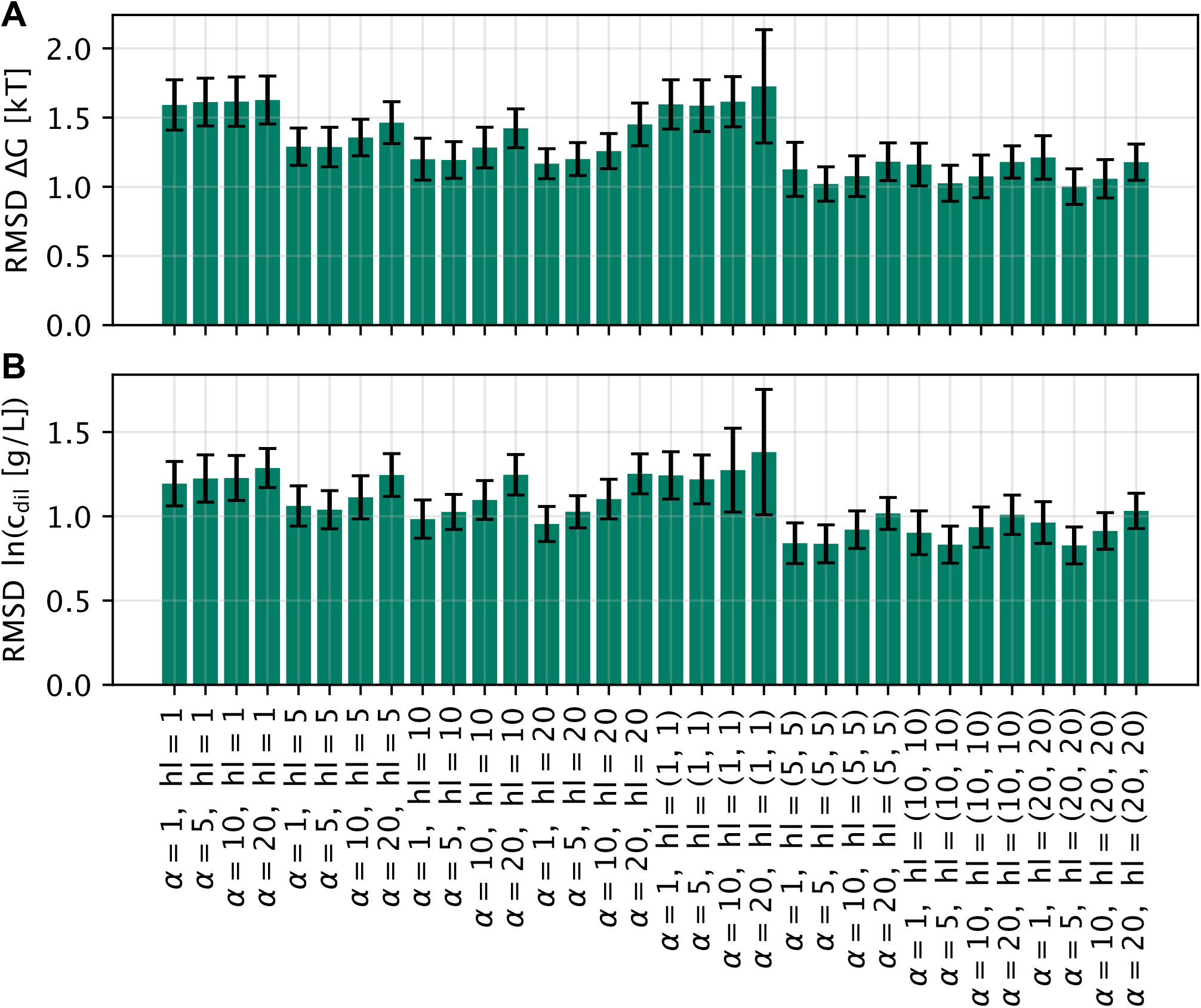
Hyperparameter search for the regularization term, *α*, and the hidden layer architecture (‘hl’) for (A) the Δ*G* and (B) the saturation concentration model. Both models have optimal parameters *α* = 5 and hl = (10, 10).

**Figure S3:**
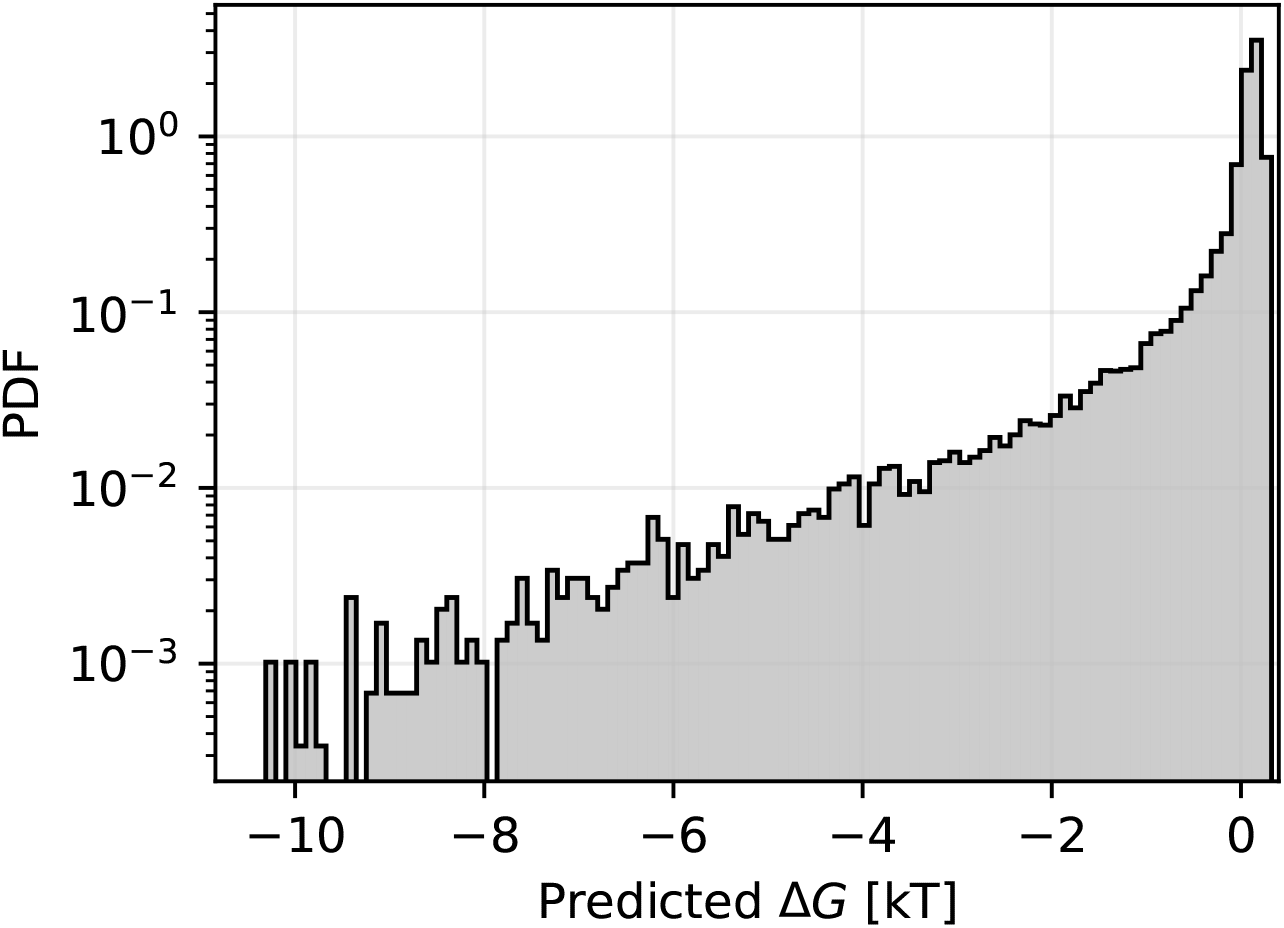
Histograms of the Δ*G* distribution for the IDRome. We note that the dynamical range of the simulations means that sequences with Δ*G <* −10 *k*_B_*T* will have calculated values of Δ*G* ≈ −10 *k*_B_*T* and sequences that are not predicted to undergo spontaneous PS (Δ*G >* 0 *k*_B_*T*) will have Δ*G* ≈ 0 *k*_B_*T*.

**Figure S4:**
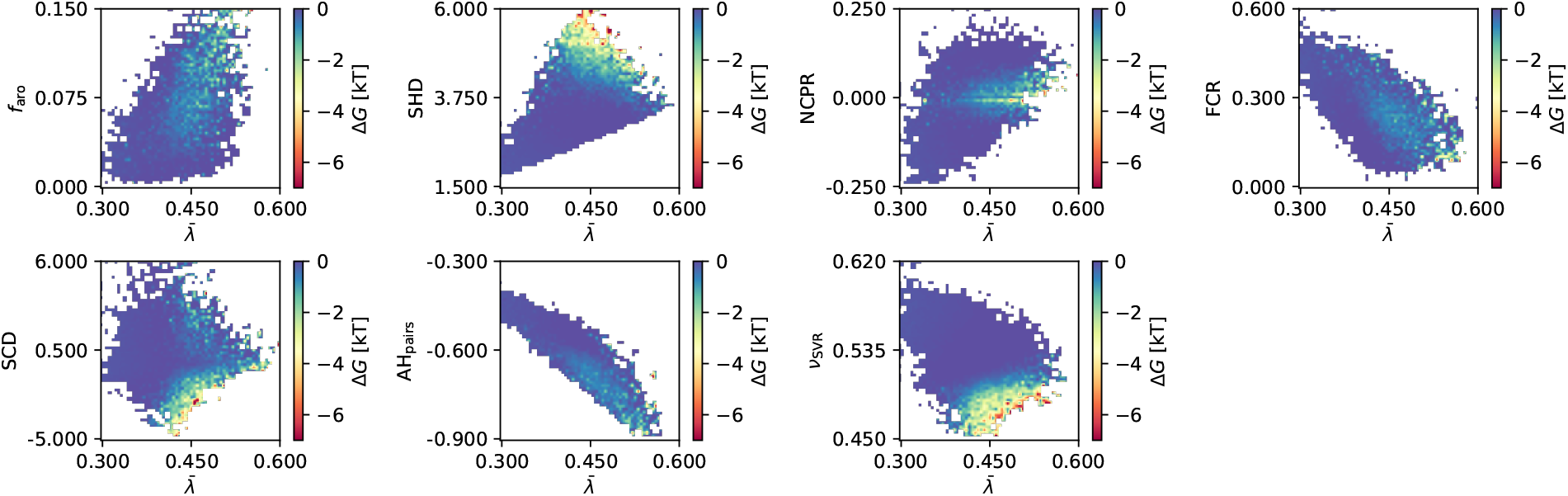
Mean values of Δ*G* for pairs of features including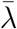. Shading from blue to red indicates increased propensity to undergo PS.

**Figure S5:**
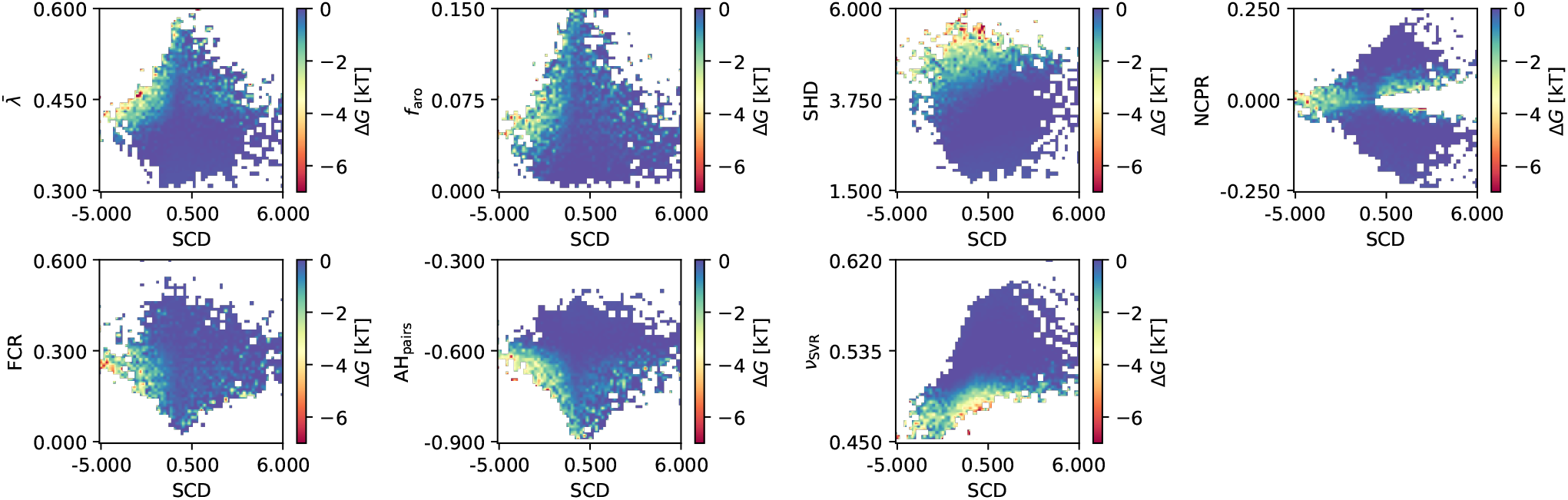
Mean values of Δ*G* for pairs of features including SCD. Shading from blue to red indicates increased propensity to undergo PS.

**Figure S6:**
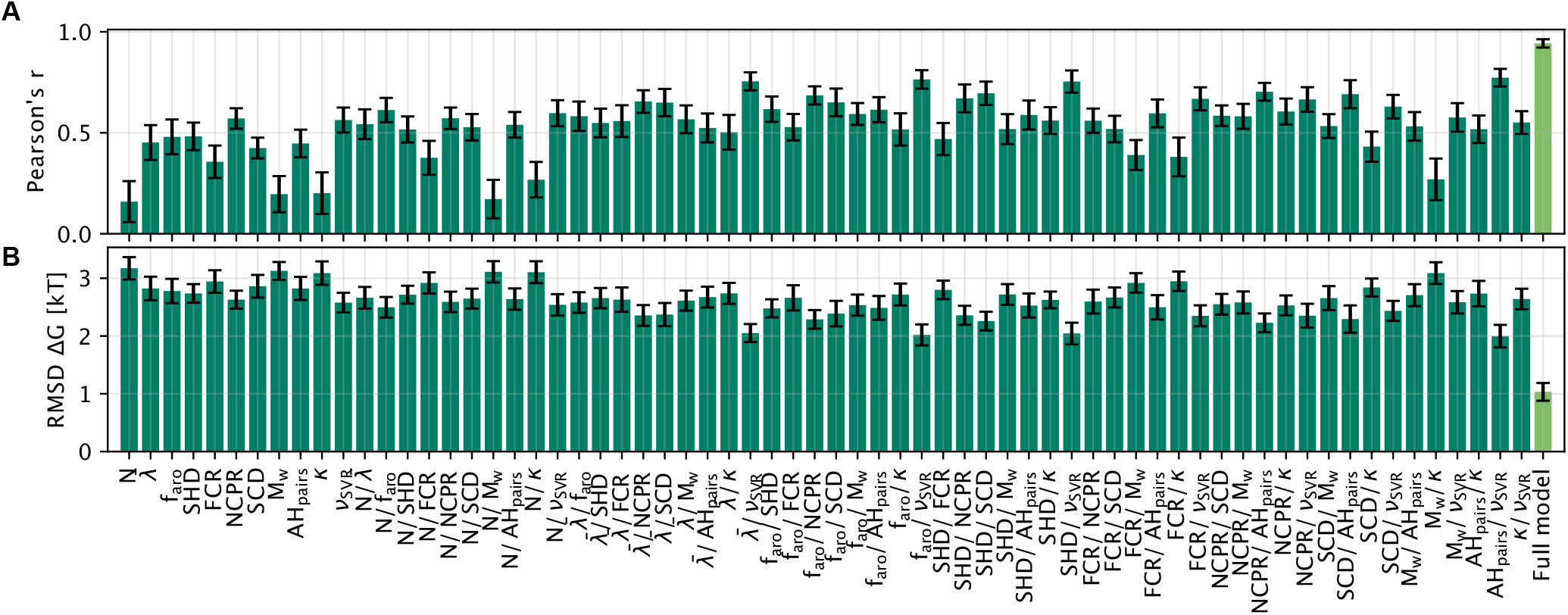
NN model for prediction of Δ*G* using combinations of up to two features as input. The model using all features listed in the main text is shown as reference. Model parameters are *α* = 5 and 2 × 10 hidden layers.

**Figure S7:**
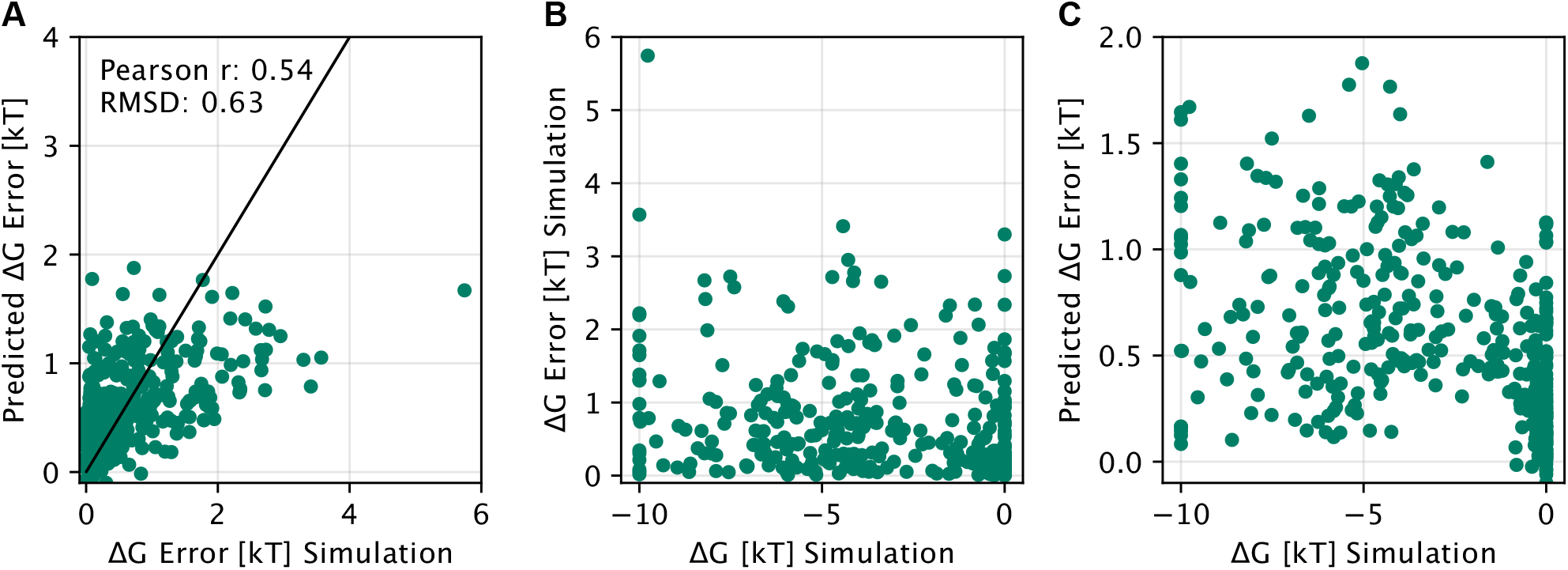
(A) Scatter plot of predicted prediction error vs. true prediction error for an error model that we trained on 389 IDRome_90_ simulations. (B) True prediction error vs. simulated Δ*G* values. (C) Predicted prediction error vs. simulated Δ*G* values.

**Figure S8:**
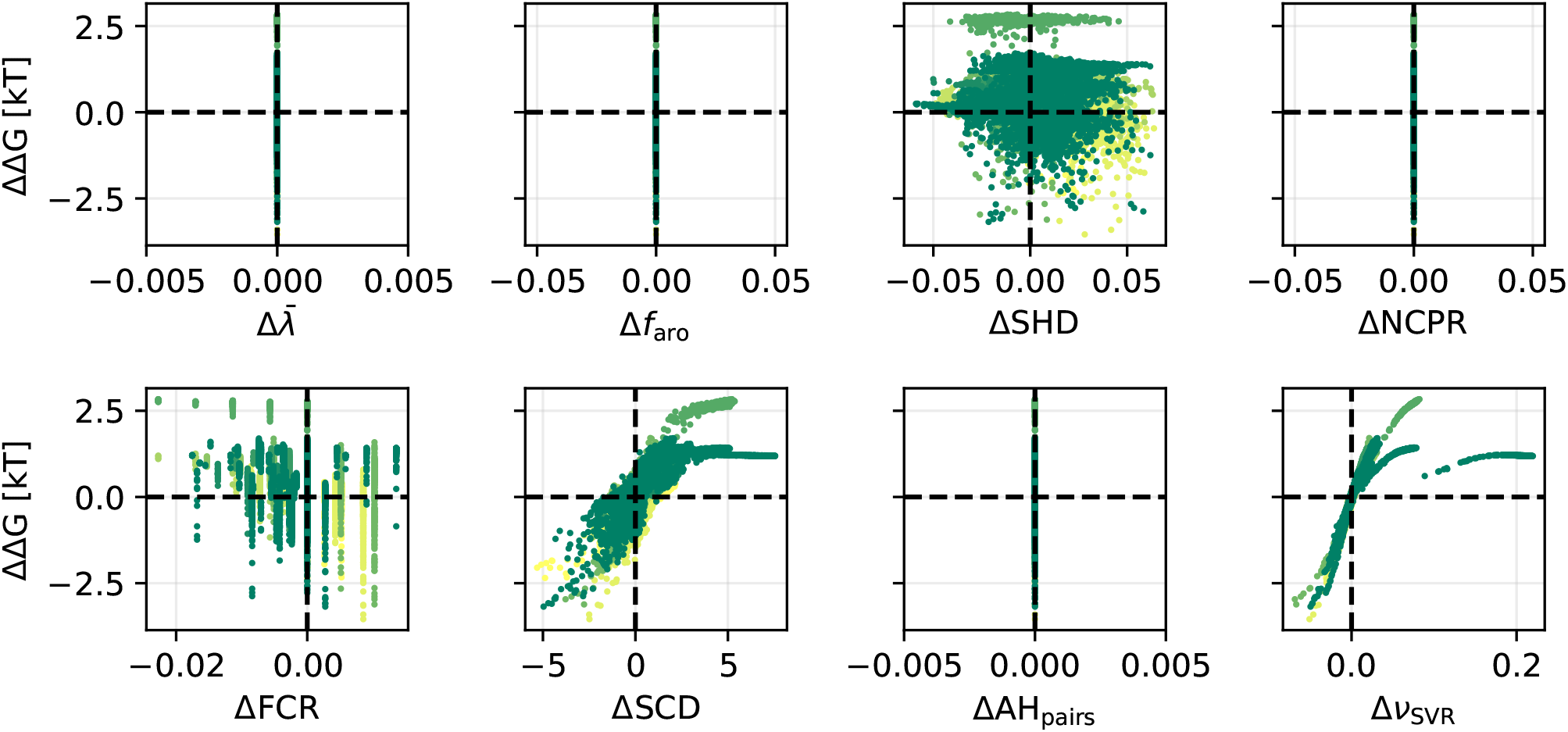
Changes to predicted PS propensity (ΔΔ*G*) for free exploration of sequences with fixed sequence composition (i.e. only allowing for swaps of amino acids). Different colours correspond to independent runs and starting points of the algorithm. The results show that changes to predicted PS propensities (ΔΔ*G*) are reflected in single chain compaction (*ν*_SVR_), and mostly driven by changes in charge patterning (ΔSCD).

**Figure S9:**
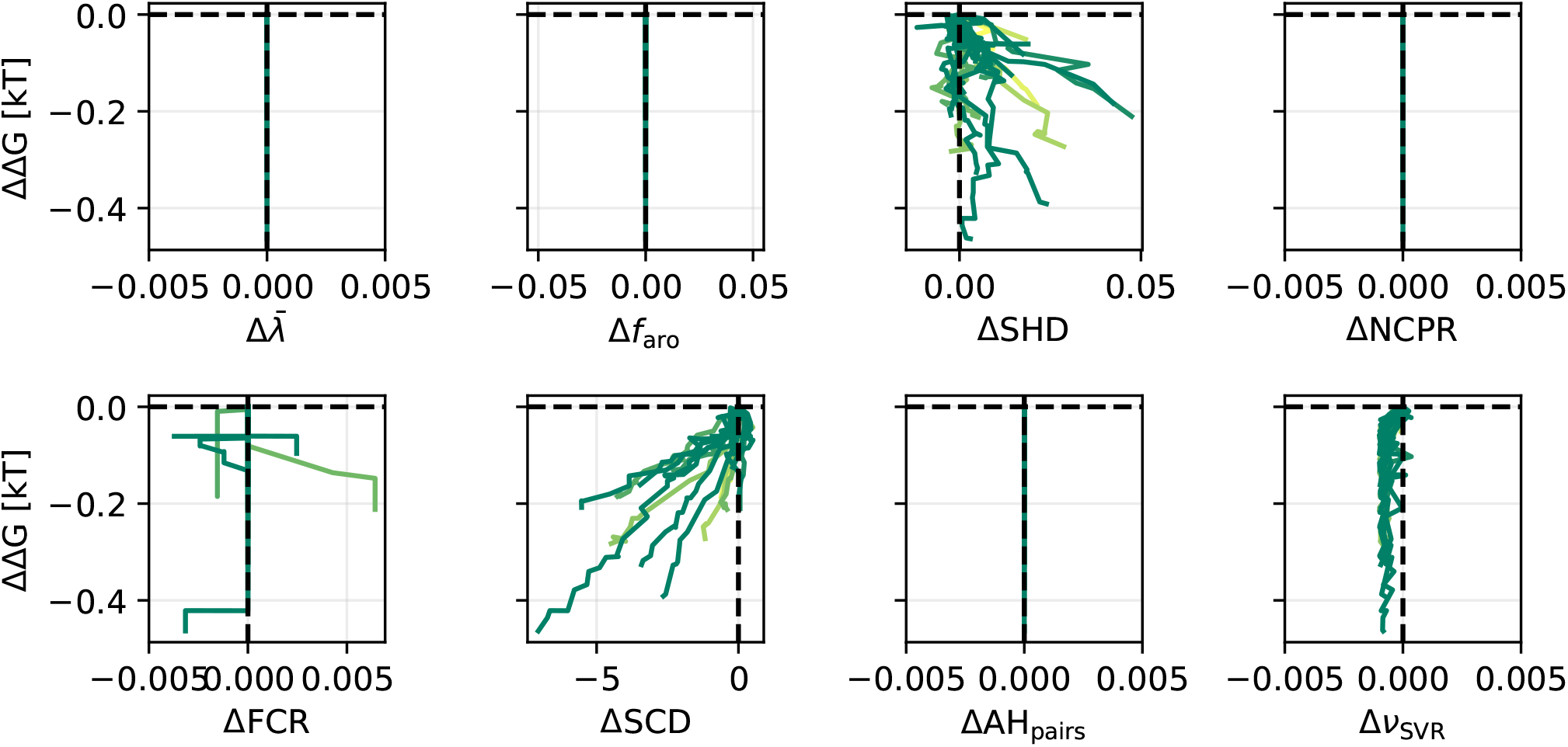
Changes to predicted PS propensity (ΔΔ*G*) for Monte-Carlo optimization towards low Δ*G*, using swap moves with *ν*_SVR_ restrained to the values of the starting sequence. Different colours correspond to independent runs of the algorithm. The results show that it is difficult to change the predicted PS propensities (ΔΔ*G*) without changing the composition and single chain compaction (*ν*_SVR_). Small changes in ΔΔ*G* are mostly driven by changes in *ν*_SVR_ within the restraint limit.

**Figure S10:**
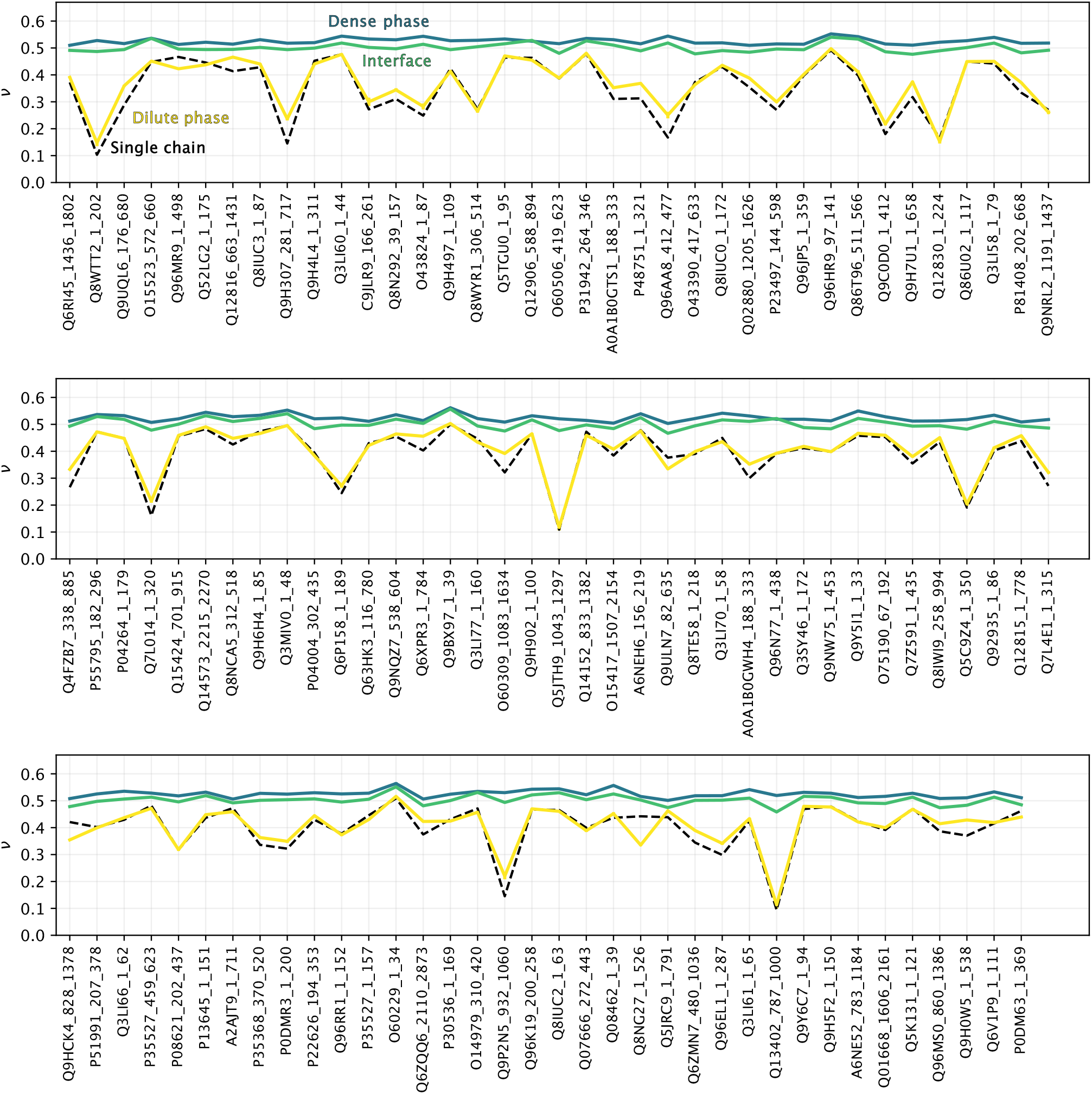
Scaling exponent *ν* from coexistence simulations of IDRome_90_ sequences simulated during the active learning protocol with −10 < Δ*G/k*_B_*T <* −4. The dilute phase, dense phase, and interface are defined based on a hyperbolic tangent fit to the concentration profile (Methods). For each frame in the trajectory, the proteins are placed in one of the three regions based on the *z*-position of the centre-of-mass of the IDR. Dashed black lines show scaling exponents from 200 ns single chain simulations with one protein in a simulation box of 25 nm x 25 nm x 25 nm. With this definition of compaction, regions and method for averaging, we find for these sequences that generally *ν*_dense_ *> ν*_interface_ *> ν*_dilute_.

**Figure S11:**
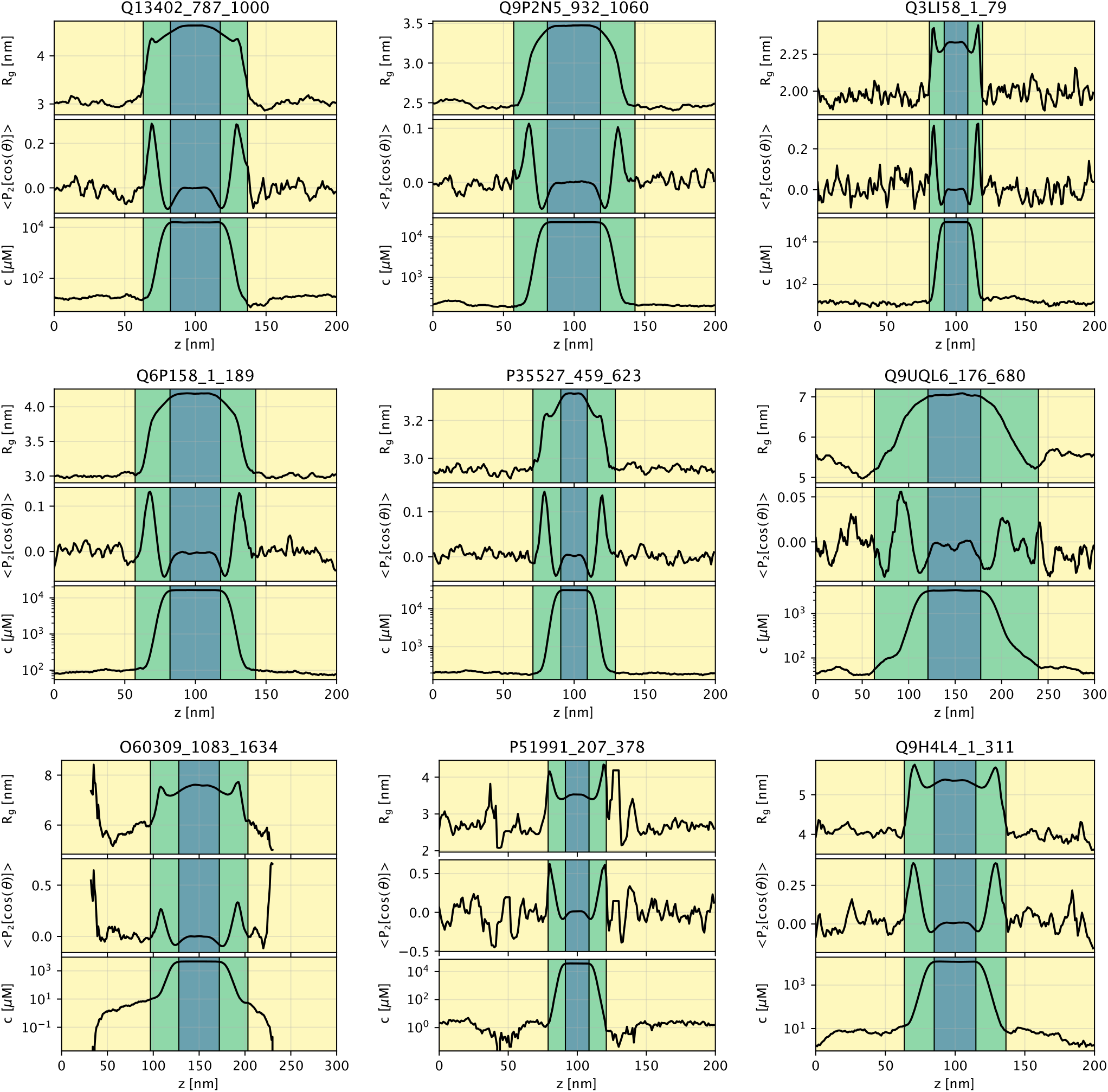
Profiles of *R*_g_, *S*_*z*_, and concentration binned along the long box edge *z* for nine examples of direct-coexistence simulations. Blue, green, and yellow shading indicate the dense phase, interface, and dilute phase, respectively.

**Figure S12:**
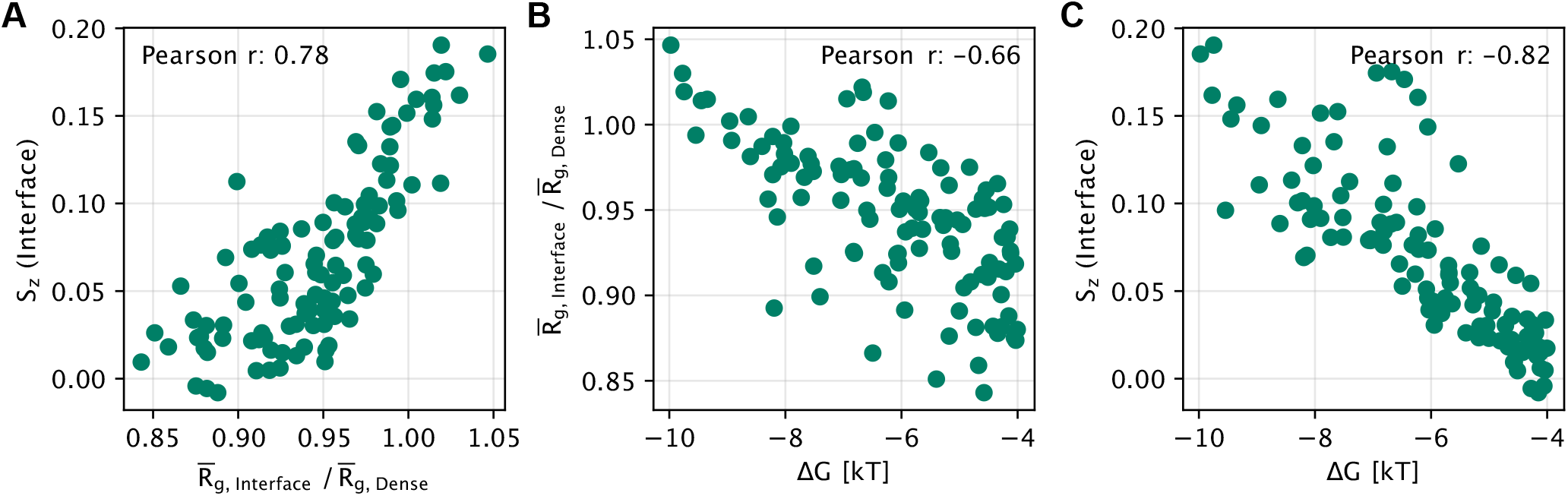
Correlation between condensate properties at the interface and transfer free energies. (A) Correlation plot of orientation order parameter *S*_*z*_ and bin-averaged 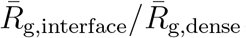. (B) Correlation plot of 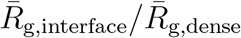 and Δ*G*. (C) Correlation plot of *S*_*z*_ and Δ*G*. The data include simulations acquired during active learning, with −10 < Δ*G/k*_B_*T <* −4.

**Figure S13:**
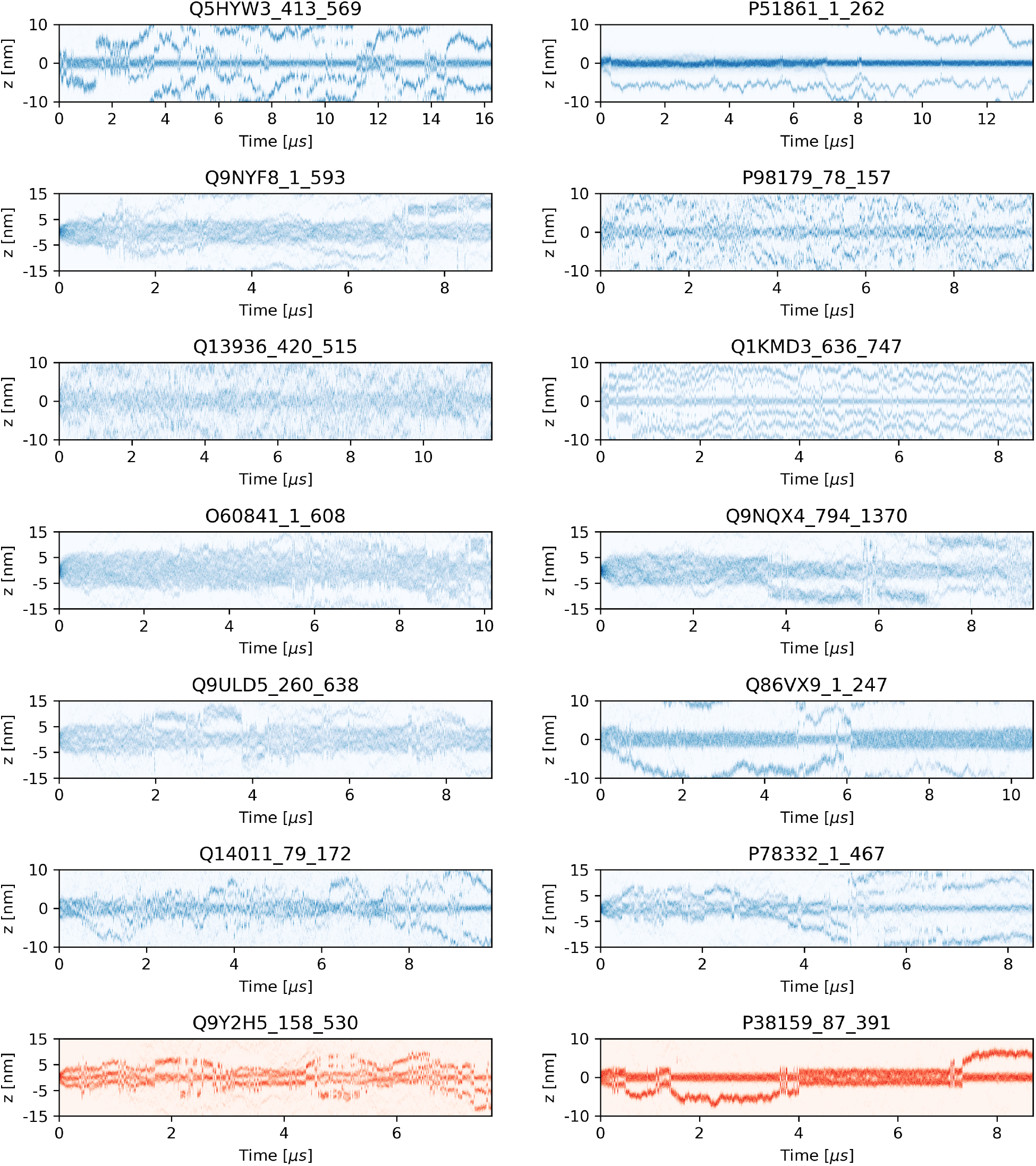
Density time traces along the *z* direction of simulation box for simulations excluded from the IDRome_90_ (blue) and IDRome_10_ (red) set.

